# Network Segregation During Episodic Memory Shows Age-Invariant Relations with Memory Performance From 7 to 82 Years

**DOI:** 10.1101/2024.12.23.630050

**Authors:** Håkon Grydeland, Markus H. Sneve, James M. Roe, Liisa Raud, Hedda T. Ness, Line Folvik, Inge Amlien, Oliver M. Geier, Øystein Sørensen, Didac Vidal-Pineiro, Kristine B. Walhovd, Anders M. Fjell

**Affiliations:** Center for Lifespan Changes in Brain and Cognition, Department of Psychology, University of Oslo, 0317 Oslo, Norway Introduction; Department of Diagnostic Physics, Division of Radiology and Nuclear Medicine, Oslo University Hospital, Oslo, Norway; Department of Radiology and Nuclear Medicine, University of Oslo, 0317 Oslo, Norway

**Keywords:** Episodic memory, functional connectivity, networks, segregation, fMRI, dedifferentiation

## Abstract

Lower episodic memory capability, as seen in development and aging compared with younger adulthood, may partly depend on lower brain network segregation. Here, our objective was twofold: (1) test this hypothesis using within- and between-network functional connectivity (FC) during episodic memory encoding and retrieval, in two independent samples (n=734, age 7-82 years). (2) Assess associations with age and the ability to predict memory comparing task-general FC and memory-modulated FC. In a multiverse-inspired approach, we performed tests across multiple analytic choices. Results showed that relationships differed based on these analytic choices, were often weak, and mainly present in the cohort with the most data. Significant relationships indicated that (i) memory-modulated FC predicted memory performance and associated with memory in an age-invariant manner. (ii) In line with the so-called neural dedifferentiation view, task-general FC showed lower segregation with higher age in adults which was associated with worse memory performance. In development, although there were only weak signs of a neural differentiation, that is, gradually higher segregation with higher age, we observed similar lower segregation-worse memory relationships. This age-invariant relationships between FC and episodic memory suggest that network segregation is pivotal for memory across the healthy lifespan.

**Declarations of interest:** none.

**Highlights:**

- Within- and between network functional connectivity predict memory performance.
- Multiverse-inspired analyses showed varying results depending on analytic choices.
- Memory associations particularly in the cohort with most data were age-invariant across the lifespan.
- Dedifferentiation might be better characterized as degrees of differentiation.

## 1. Introduction

A proposed factor behind cognitive decline in aging is neural dedifferentiation (Koen and Rugg, 2019). Neural dedifferentiation refers to neural processing becoming less selective in older adults, putatively compromising processing fidelity and efficiency (Li et al., 2001).

Using functional magnetic resonance imaging (fMRI), neural dedifferentiation has been investigated at three levels: individual items, perceptual categories, and functional brain networks (Koen et al., 2020). Findings at the item- and category levels suggests that (i) neural dedifferentiation may be particularly important for explaining age-related decline in episodic memory (Koen et al., 2019), and (ii) that the relationship between dedifferentiation and memory do not differ in strength as a function of age, that is, it is age-invariant (Koen et al., 2020). However, at the level of functional networks, few studies have examined dedifferentiation and memory performance, particularly while measuring network dedifferentiation during memory processing, likely more closely related to episodic memory performance (Greene et al., 2018; Greene et al., 2023). It is therefore unknown whether also network-level dedifferentiation demonstrate age-invariant relationships with memory performance across the adult lifespan. Such a relationship would suggest that higher network selectivity accompanies better memory performance not only in aging, but also earlier in the lifespan (Rugg, 2017).

Indeed, the term dedifferentiation implies a preceding differentiation. Yet, we lack studies contrasting network-level degree of differentiation and cognitive performance in both development and aging. In line the age differentiation-dedifferentiation hypothesis (Li et al., 2004; Tucker-Drob, 2009), we would expect network differentiation in development, linked with the concurrent increase in episodic memory capabilities (Ngo et al., 2018), to precede network dedifferentiation in aging, in turn linked with memory decline (Park and Reuter-Lorenz, 2009). In other words, an age-invariant, positive lifespan pattern, with higher degree of network differentiation associating with higher memory capabilities.

Demonstrating an age-invariant, lifespan pattern would be important for at least two reasons. First, as mentioned, it would suggest a prominent, lifelong role for the degree of network-level differentiation for memory and potentially other cognitive processes. Second, such findings would lend confidence in network-level degree of differentiation being an important biomarker not only in cognitive aging, but also in cognitive development. Finally, lack of support, as manifested for lifespan differentiation-dedifferentiation of cognitive abilities (Tucker-Drob, 2009), would also be of important theoretical significance, given the implications for theories of neurocognitive development and aging.

What is the relationship between functional networks and episodic memory through the lifespan? Network-level dedifferentiation using functional connectivity (FC) manifests in older age as lower connectivity within a network, and higher connectivity between networks, that is, lower network segregation. Below, we use this segregation term, which is more theoretically neutral, and refers to the neuronal processing carried out among functionally related regions arranged within networks or modules (Chan et al., 2014; Sporns, 2013).

Previous studies using resting-state fMRI (rs-fMRI), have reported age-related differences in segregation, that is, lower within-network FC and higher between-network FC at older ages (Andrews-Hanna et al., 2007; Cassady et al., 2020; Cassady et al., 2019; Han et al., 2018; Madden et al., 2020; Meunier et al., 2009; Setton et al., 2022; Stumme et al., 2020; Zonneveld et al., 2019). This lower segregation has been linked with lower memory performance (Chan et al., 2014; Stumme et al., 2020; Varangis et al., 2019), albeit not uniformly (Cassady et al., 2021; Chong et al., 2019). These findings are in line with the dedifferentiation view discussed above (Koen and Rugg, 2019). The strength of these associations between rs-fMRI-based FC and out-of-scanner memory was not tested, and hence unclear whether they were age-invariant (Koen et al., 2020). In development, a lifespan rs-fMRI study reported mainly lower segregation with higher age from 7 years (Betzel et al., 2014), while Petrican et al. (2017), in subsamples below 20 years of age, did not observe associations between episodic memory and within-network FC.

In adults, partly opposite results stem from two studies with networks derived from task-fMRI data. Both Grady et al. (2016) and Monge et al. (2018) found higher between-network FC in older adults compared with younger adults, for fronto-parietal and medial temporal lobe FC, respectively. Grady et. al. found that this higher between-network FC was related to better post-scanning associative memory performance, in line with Cassady et al. (2021), only in older adults, and hence this relationship was age-variant. In contrast, Monge et al. found that the higher between-network FC was related to better source judgement across both the younger and older age groups, that is, the memory relation was age-invariant. These findings contrast with the dedifferentiation view of worse memory with less segregation, as measured by higher between-network FC. Instead, according to Monge et al. (2018), despite their age-invariant memory relation, their finding fits a view of less segregation reflecting compensatory processes that are beneficial for cognition.

Finally, how are relationships affected by different ways of analyzing FC? Analysis of FC can be used to reveal several key aspects of brain activity, and different analytical approaches might influence the sensitivity of task-fMRI to behavioral performance differences (Greene et al., 2018). Traditionally, FC has been used to capture "intrinsic" BOLD fluctuations, reflecting ongoing, spontaneous, and possible state-dependent brain activity.

This included the two studies mentioned above: Grady et al. (2016) regressed out specific task effects during item encoding. This approach yields connectivity patterns related to the ongoing connectivity as well as residual, general task effects. Monge et al. (2018) used a correlational psycho-physiological interactions (cPPI) approach (Fornito et al., 2012), which, when derived from one task event like source memory, in effect also captures condition-unspecific connectivity similar to rs-fMRI FC (Raud et al., 2023). Hence, such approaches probe brain characteristics that more strongly reflect a condition-unspecific task-state and background task FC (here, for simplicity, termed “task-general FC”).

Additionally, using a generalized PPI (gPPI) approach (McLaren et al., 2012), FC can quantify task- and even condition-specific co-fluctuations by isolating these from the condition-unspecific activity, potentially highlighting how brain networks adjust in response to specific task demands. Hence, here, gPPI likely yields condition-specific FC more strongly modulated by memory processes. Whether such memory-modulated FC shows segregation differences across the lifespan and how it relates with memory has not previously been investigated.

Here, we investigated these knowledge lacunas using task-fMRI during encoding and retrieval of episodic information, in two independent samples undergoing similar memory tasks, in data covering most of the lifespan (total n=734, age 7-82 years). As measures of segregation, we used within- and between-network FC. We compared condition-unspecific, task-general FC and memory-modulated FC, at both encoding and retrieval, and assessed relations with age and in-scanner memory performance. In addition to replication across the two datasets, we also tested the generalizability of segregation-memory relations using leave-one-out cross-validation (LOO-CV) (Rosenberg and Finn, 2022). To assess the potential diversity of results arising from different analytical choices, we performed the analyses across several approaches in a multiverse analysis-inspired manner, and present results using specification curves (Simonsohn et al., 2015, 2020).

We hypothesized less network segregation with older age, that is, lower within-network FC and higher between-network FC for task-general FC, for most networks except visual networks (Zonneveld et al., 2019). Based on Betzel et al. (2014), we expected a similar pattern in development. Second, we tested whether we would observe different patterns for memory-modulated FC. Third, in the task-general FC analyses, as less segregation was expected with higher age, we hypothesized that less segregation would relate to worse source memory performance. Finally, for memory-modulated FC, we tested whether relations with memory would show a different pattern.

## 2. Material and methods

### 2.1 Participants

All participants ≥12 years gave written, informed consent. All participants <12 years gave oral informed consent and, for all participants <18 years, written informed consent was obtained from their legal guardians. Ethical approval was obtained by the Regional Committees for Medical and Health Research Ethics in Norway.

For the purpose of current analysis, four samples were constructed from two independent, cross-sectional samples performing two different memory tasks, associated with the ERC funded projects Constructive Memory (https://cordis.europa.eu/project/id/283634) (Raud et al., 2023; Vidal-Pineiro et al., 2021), analyzed for 544 participants with fMRI data, with a continuous age distribution from 6 to 82 years, and the AgeConsolidate (https://cordis.europa.eu/project/id/725025) (Ness et al., 2022), with a total of 245 participants, distributed across three age groups, from 11 to 81 years. The youngest participants in the Acon sample (< 20 years) went through a shortened study protocol without in-scanner fMRI retrieval (see below).

Due to the age distribution of the data, previous influential adult lifespan studies focusing on the age span from 20 to 80 years (Chan et al., 2014), and potential non-linearity across the lifespan complicating comparisons across tests, we created four sample, two adult samples and two developmental samples. Specifically, we also divided the Constructive Memory sample in (i) a developmental (Dev1) and (ii) an adult (Adult1) subsample (**Table 1**). Similarly, we divided the AgeConsolidate sample into two: (i) a developmental (Dev2) subsample (10–16 years), with only encoding in scanner, and (ii) an adult (Adult2) subsample, by collapsing younger (20–44 years) and older (60-81 years) adults into one group.

**Table 1.**
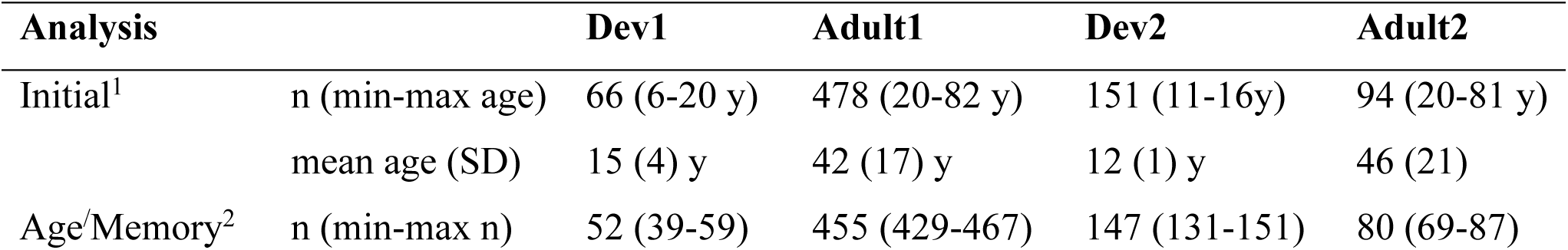
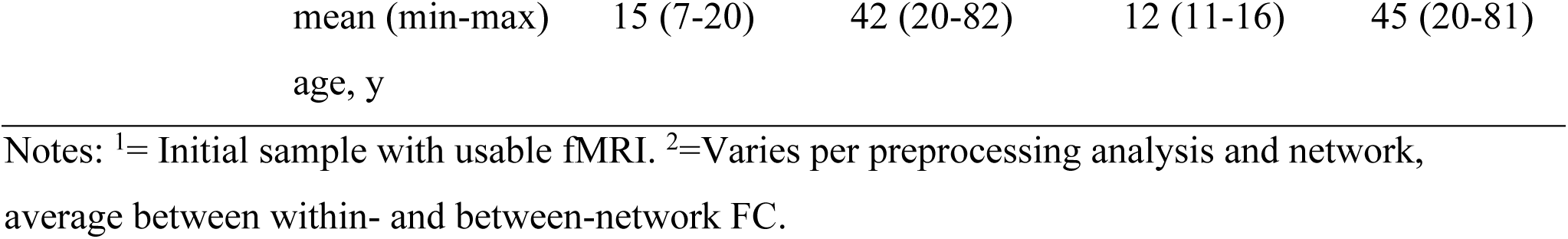
Sample size and age.

The age distribution in the subsamples were not normally distributed (P < 0.05). For the regression models, it is the residuals that should be normally distributed for proper inferences, although, for larger samples, there should be little issue with deviations from normality. For the Dev1, with sample size < 50, we tested the normality of the residuals in each model. We only found a minority of deviations from normality (1% or below for TGFC, while MMFC showed 20% for within- and 2% for between-network FC). In instances where residuals were non-normal, we ran a robust regression and compared the t values. The comparisons showed that for these few cases, showed relatively little differences in t values, which were > 2 in all but 3 instances (1.9, 1.9 and 1.7).

Exclusion criteria were MRI contraindications, history of injury, disease or psychoactive drug use affecting central nervous system function, including clinically significant stroke, psychiatric illness, serious head injury, untreated hypertension, and diabetes, as well as MRI contraindications, and, for the Adult2 sample, sleeping disorders.

We discarded observations due to partially missing and/or deviant neuropsychological data. On the Mini-Mental State Examination (Folstein et al., 1975), participants above 40 years of age scored ≥26, except 2 Adult1 participants aged 42 and 80 years scoring 25, which were included due to non-deviant memory scores. Participants had full-scale IQ above 85 on the Wechsler Abbreviated Scale of Intelligence (Wechsler, 1999). Hence, the samples were without signs of cognitive impairments. All participants who completed the Beck Depression Inventory (BDI) scored ≤17, except two Adult2 participants aged 27 and 28 scoring 21 and 19, respectively, also included due to non-deviant memory scores. Participants 60 years or older in the Adult2 sample (missing for 12 participants, two of which had a BDI score of 5) performed the Geriatric Depression Scale (Yesavage et al., 1982), and scored ≤ 7.

### 2.2 fMRI memory tasks

The Adult1 and Dev1 subsamples performed an incidental memory encoding task, as described previously (Sneve et al., 2015), followed by a surprise retrieval phase after ∼90 minutes. Shortly, during encoding, we presented 100 black and white line drawings depicting everyday objects. In the first half, at the presentation of the drawing, the participants heard the question ‘Can you eat it?’, and in the second half, the question ‘Can you lift it?’, or vice versa. The task was to answer the question by a button press indicating ‘Yes’ or ‘No’ (button-response mapping counterbalanced across participants). At retrieval, we presented all 100 items from encoding intermixed with 100 new items. For each item, the task was to answer the question ‘Have you seen this item before’. If the participant answered ‘Yes’, a second question was asked: ‘Can you remember what you were supposed to do with the item?’. If the participants answered ‘Yes’, a third question was asked: ‘Were you supposed to eat it or lift it?’, and the participants had to choose between answers ‘Eat’, and ‘Lift’. In case of ‘No’ answer, the trial ended.

For this Adult1 and Dev1 memory analysis, a source recollection index was calculated as a percentage of correct associations from all items. Approximate guessing rate for each participant was estimated by the number of incorrect answers to ‘Eat’/’Lift’, retrieval questions, as the participants had indicated that they remembered the association, but then chose the wrong answer. These were subtracted from the number of correct answers before calculating the recollection index to correct for guessing (Raud et al., 2023; Sneve et al., 2015; Vidal-Pineiro et al., 2021). Task presentation was controlled with E-Prime 2.0.

The Adult2 and Dev2 samples performed an instructed memory encoding phase, an intermediate forced choice retrieval phase directly after encoding, and a recollection retrieval phase 12 h after encoding (Ness et al., 2022; Raud et al., 2023). The encoding and 12-hours retrieval were performed in the MRI scanner for the Adult2 subsample. Dev2 participants did not return for the 12-hours retrieval sessions, thus only encoding fMRI data is available for Dev2. For Adult2, most participants completed the procedure twice, over two separate visits, separated by a minimum of 6 days. The time of the day for encoding and the forced choice memory test was counterbalanced within-subjects, with one encoding and test session completed in the evening in one visit and the other encoding and test session completed in the morning during the other visit. Task data from both visits were collapsed to calculate memory scores for the Adults2 sample. The trials in the encoding and final retrieval phase were separated into 2 and 3 runs, respectively, with 64 trials for each run. During the encoding phase, participants were presented with 128 item- and face or place pairs per visit (drawn from a pool of 256 items, 8 faces, 8 places), accompanied by an auditory recording that repeated the Norwegian word for each item three times. The items consisted of real-life images of non-animate everyday items. For a single session, 4 different faces and places were used. Participants were instructed to imagine an interaction between the item and the face or place while presented on the screen, and afterwards to rate the vividness of their imagination on a scale from 1 to 4, in which 1 was “not vivid at all” and 4 was “very vivid”. They were informed that the item-face/place associations would be tested in later sessions. After the encoding phase, all participants completed a self-paced forced-choice retrieval task with all 128 trials outside of the scanner (64 trials for Dev2). In each trial, they were presented simultaneously with the image and the recording of one of the learned items, together with all 4 faces and 4 places used in the session. Their task was to indicate by a button press, whether the item had been paired with a face or a place during the encoding phase. For Dev2 participants, the memory score was calculated as the number of correct answers (divided by 64, i.e., the number of trials). The Adult2 subsample returned for a retrieval phase 12 h later. Here, they were presented with 128 learned items and 64 new items (both visual and auditory) in a pseudorandomized order. The participants had to a press a button choosing between four options: (O1) they had seen the item before and it was associated with a face, (O2) they had seen the item before and it was associated with a place, (O3) they had seen the item before, but could not remember whether it was associated with a face or place, (O4) they had not seen this item before. In the memory analyses, correct answers to O1 and O2 were considered source recollection for Adult2. The recollection rates were corrected for guessing by subtracting the number of incorrect associations from the number of correct associations.

Task presentation was controlled with Psychtoolbox 3. In both studies, the inter-stimulus interval was determined using optseq2 (https://surfer.nmr.mgh.harvard.edu/optseq/), and a fixation cross remained on the screen until the beginning of the next trial (for details, please see Sneve et al. 2015, and Ness et al. 2022). Memory performance for each subsample is presented in **Fig. 1** (see details in Section 3.1 below).

**Figure 1.**
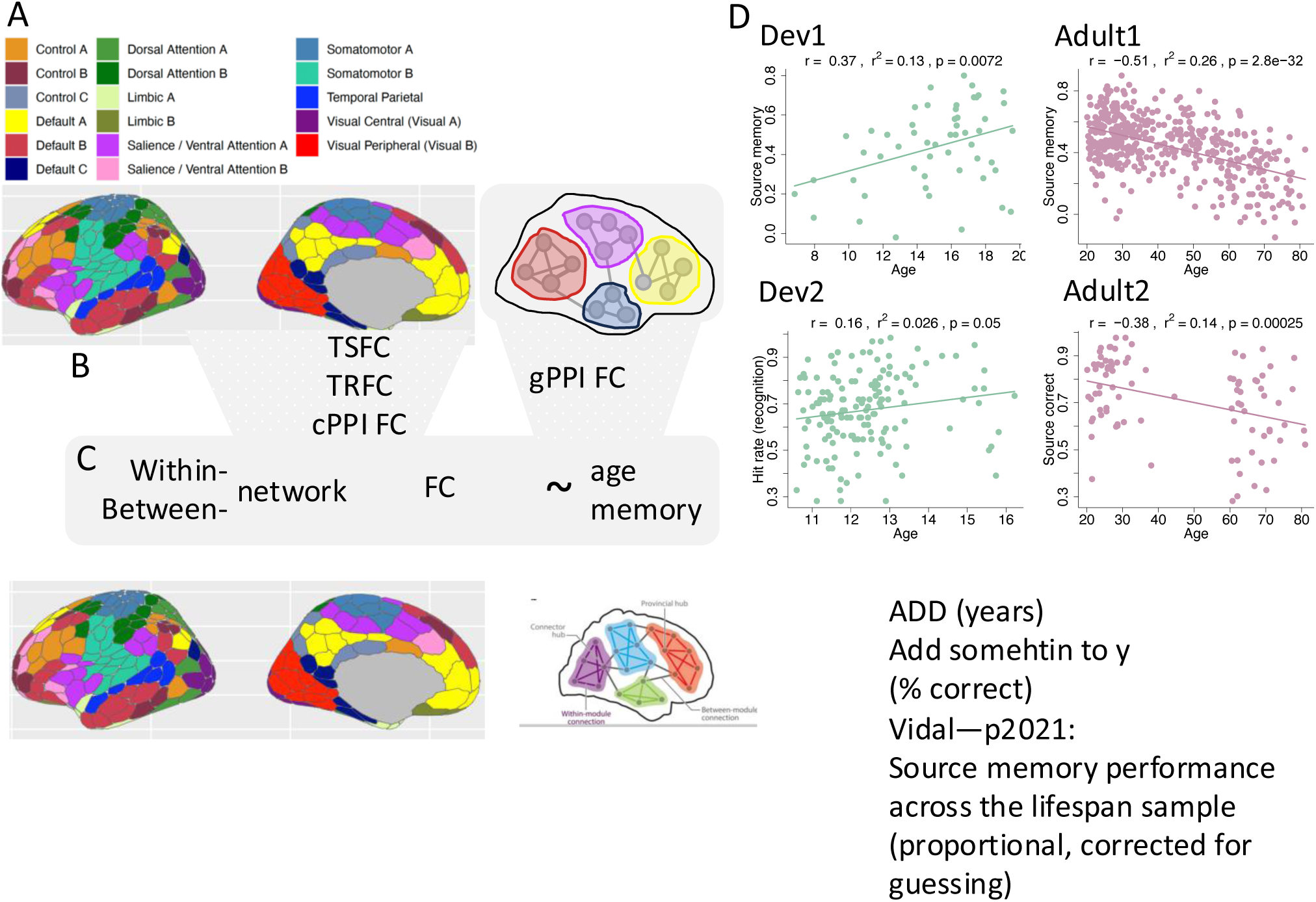
Study design and memory performance. **A**. The Schaefer 400 region atlas and their network affiliation, used to obtain (**B**) the three different FC measures: TSFC=task-state FC, BGFC=background FC, cPPI=correlation psychophysiological interactions [PPI]. For generalized PPI (gPPI), we used the data to calculate modules/networks. **C**. Within- and between network values from these four kinds of FC were then associated with age and memory performance. **D**. Memory performance with age in the four subsamples. Atlas plotted using *ggseg* (Mowinckel and Vidal-Piñeiro, 2019)

### 2.3 MRI acquisition

Adult1 and Dev1 performed the memory task on a 3T Skyra scanner (Siemens Medical Solutions, Germany) at Rikshospitalet, Oslo University Hospital, using a 20-channel head-neck coil. The parameters were equivalent for encoding and retrieval phases. For each run, 43 transversally oriented slices were measured using a blood oxygen level–dependent (BOLD)- sensitive T2*-weighted EPI sequence (TR = 2390 ms, TE = 30 ms, flip angle = 90⁰; voxel size 3 × 3 × 3 mm^3^; FOV = 224×224 mm^2^; interleaved acquisition; GRAPPA factor = 2).

Each encoding run consisted of 131 volumes, while the number of volumes during retrieval was dependent on participants’ responses (207 volumes on average). Three dummy volumes were collected at the start of each fMRI run to avoid saturation effects in the analyzed data. Additionally, a standard double-echo gradient-echo field map sequence was acquired for distortion correction of the EPI images. Anatomical T1-weighted MPRAGE images, consisting of 176 sagittal slices, were obtained using a turbo field echo pulse sequence (TR = 2300 ms, TE = 2.98 ms, flip angle = 8°, voxel size = 1 × 1 × 1 mm^3^, FOV = 256×256 mm^2^).

Adult2 and Dev2 participants performed the memory task on a Prisma 3T scanner (Siemens Medical Solutions, Germany) at Rikshospitalet, Oslo University Hospital, using a 32-channel head coil. The parameters were equivalent for encoding and retrieval task phases. For each run, 56 transversally oriented slices were measured using a BOLD-sensitive T2*- weighted EPI sequence (TR = 1000 ms; TE = 30 ms; flip angle = 63°; matrix = 90×90; voxel size 2.5 × 2.5 × 2.5 mm; FOV = 225×225 mm; ascending interleaved acquisition; multiband factor = 4, phase encoding direction = AP). Each encoding and retrieval run consisted of 730 volumes. Six dummy volumes were collected at the start of each fMRI run to avoid saturation effects in the analyzed data. Sufficient T1 attenuation was confirmed following preprocessing. Additional spin-echo field map sequences with opposing phase encoding directions (anterior-posterior and posterior-anterior) were acquired for distortion correction of the EPI images.

Anatomical T1-weighted MPRAGE images, consisting of 208 sagittal slices, were obtained using a turbo field echo pulse slices (TR = 2400 ms; TE = 2.22 ms; TI = 1000 ms; flip angle = 8°; matrix = 300×320×208; voxel size = 0.8 × 0.8 × 0.8 mm; FOV = 240×256 mm). T2-weighted SPACE images, consisting of 208 sagittal slices, (TR = 3200 ms; TE = 5.63 ms; matrix = 320×300×208; voxel size = 0.8 × 0.8 × 0.8 mm; FOV= 256×240 mm) were also obtained.

### 2.4 fMRI preprocessing

The fMRI data was preprocessed using FMRIPREP version 1.5.3 2 (Esteban et al., 2020; Esteban et al., 2019) and Python 3.8.2. The detailed pipeline has been described previously (Ness et al., 2022; Raud et al., 2023). Shortly, the pipeline included BIDS-conversion, intensity nonuniformity correction, skull-stripping, susceptibility distortions correction, motion correction, co-registration with the anatomical reference using boundary-based registration with 6 degrees of freedom, and slice-timing correction. Distortion correction was performed using a custom implementation (https://github.com/markushs/sdcflows/tree/topup_mod) of the TOPUP technique (Andersson et al., 2003). We applied the ICA-AROMA method (Pruim et al., 2015) for denoising, incorporating mean white matter and cerebrospinal fluid timeseries as nuisance regressors, along with six motion parameters derived from rigid body estimation. Concurrently, we executed detrending and high-pass filtering at 0.008 Hz using a Butterworth 5th order filter, ensuring orthogonality to the nuisance regressors (Hallquist et al., 2013). In FC analyses, global signal regression (GSR), has been efficient at removing signal contributions from non-neuronal physiological processes like respiration (Birn et al., 2006; Power et al., 2017b) and alleviate motion-related confounds (Ciric et al., 2017). On the other hand, GSR removes both the global physiological noise and some of the neural signal in a network-specific way, from both non-cognitive and so-called task-positive regions (with less impact on so-called task-negative regions) (Glasser et al., 2018). Hence, here we do all analyses once with GSR, and once without. In-scanner movement tends to correlate with FC estimates despite rigorous data cleaning and de-noising (Ciric et al., 2017). We ran all analyses with average framewise displacement (FD) as a covariate, and only including participants with FD < 0.3.

### 2.5 Functional connectivity and network construction

We divided the cortex into 400 regions on each participant’s native reconstructed surface (Schaefer et al., 2018) (**Fig. 1**). This parcellation stems from resting-state fMRI data. For each participant, the atlas was transformed from group template space to subject native space.

Different types of FC can be estimated from the BOLD signal during a task. Here, the FC during the encoding and retrieval tasks was calculated in a total of 4 ways (**Fig. 1** and **Table 2**): 3 ways for what we call the condition-unspecific, task-general FC (TGFC), and 1 for what we call the memory-modulated FC (MMFC). That is, for TGFC, the task data during encoding and retrieval, was processed by (1) treating the data as resting-state timeseries. Here, we refer to this as task-state FC (TSFC). (2) Regressing out the task activations and correlate the residuals (see details below). We refer to this as background FC (BGFC). (3) Using correlational PPI (cPPI). Note, cPPI is usually included as task-modulated FC, but given our previous findings of large similarity between rs-fMRI FC and cPPI FC (Raud et al., 2023), we here include cPPI as TGFC (see details below). For the MMFC, we used (4) generalized PPI (gPPI). In the 3 latter approaches, the analyses were run only on trials with correct source memory performance at retrieval and the corresponding trials at encoding (for Dev2, only encoding). In Adult2, these trials taken from the intermediate forced choice retrieval phase directly after encoding.

**Table 2.**
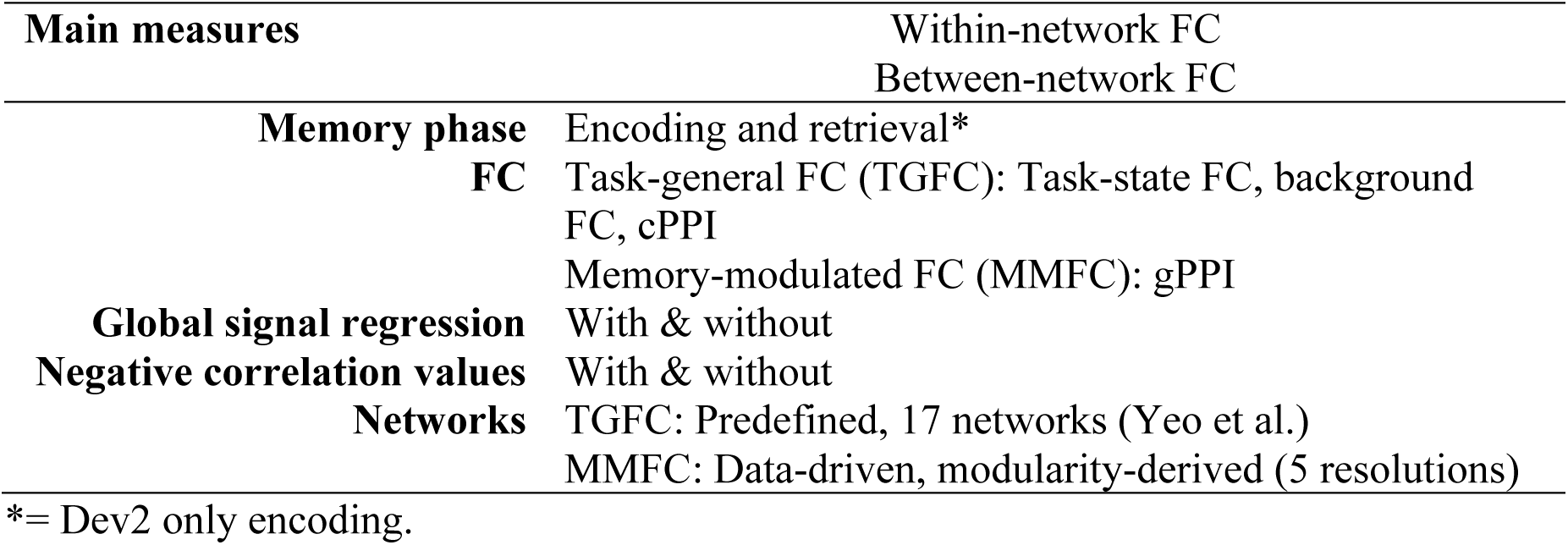
Multiverse-inspired analyses. Within- and between network FC measured separately for different memory phases, types of FC, processing steps, and networks.

More specifically, for the TGFC (1), we treated the data as resting-state timeseries, ignoring task information, and performed Pearson’s correlation between timeseries of the 400 regions The BGFC (2) was based on the time-series residualized for task events, which is designed to capture network reorganization due to changes in sustained cognitive states (Fair et al., 2007; Norman-Haignere et al., 2012). The mean evoked responses of all task events were fitted using finite impulse response models (i.e., stick regressors covering ∼25 s post-event onset) and then removed from the time series using general linear models. The number of task events modeled and removed ranged from three to eight (see **Supplemental Table** [**STable**] **1** for specifics across cohorts and memory processes). The residualized time-courses between each pair of cortical regions were then correlated, forming the so-called residualized estimates of FC.

The cPPI analysis (3) (Fornito et al., 2012) yielded symmetrical, undirected matrices reflecting the 400 regions’ FC to each other during correct source memory operations.

Briefly, the psychological timeseries were constructed as boxcar functions, reflecting 2 s (Dev1/Adult1) or 5 s (Dev2/Adult2) events from the start of the trial, as this matched the duration of the memory relevant stimulus presentation in the two experiments, respectively. The PPI terms (**STable 1**) were constructed by first deconvolving BOLD timeseries from each region (Gitelman et al., 2003), then multiplying these point-by-point with the psychological event timeseries, and finally re-transforming these to BOLD through convolution with a canonical 2-gamma HRF. Finally, the cPPI matrices were constructed by partial Pearson’s correlation analysis between the constructed PPI terms of each pair of the cortical regions, controlling for each region’s BOLD timeseries and the HRF-convolved psychological timeseries. For example, for encoding in the Adult2 cohort, the (MatLab) formula was: partialcorr(PPI:SourceROI1, PPI:SourceROI2, covariates), where the PPI:Source variables for ROI1 and ROI2 refer to the source memory condition PPI term for the seed (ROI1) and the target (ROI2) region, respectively. The covariates were the psychological variables (task design regressors for source, miss, and response), and the physiological variables (BOLD time series) from the seed and test region, respectively (see **Supplemental Figure [SFig.] 1** for a matrix representation of the same example, and **STable 1** for specifics across cohorts and memory processes). The PPI variable was calculated simultaneously for all source/hit memory trials. As such, it represents event-related connectivity contrasted to implicit baseline during the task period and is similar to the beta-series correlation technique (Di et al., 2021), and captures both condition-unspecific connectivity and task-specific connectivity. In effect, this yields FC similar to the TSFC and BGFC approaches outlined above and rs-fMRI analysis (Raud et al., 2023).

For MMFC analysis, we performed region-of-interest-based gPPI (McLaren et al., 2012) as described in Di and Biswal (2019), in the 400 regions. As for cPPI, BOLD timeseries for each region (first eigenvariate over voxels) were deconvolved and multiplied by the centered boxcar regressor representing the event in “neuronal space”. The resulting “neuronal” PPI terms were re-convolved with a HRF resulting in BOLD-level PPI predictors. Unlike cPPI, the gPPI measure of region-to-region FC is established via regression where region A’s observed BOLD timeseries gets explained by a matrix of explanatory variables consisting of a) region B’s PPI term for the psychological task of interest, b) region B’s PPI terms for other modeled task events, c) region B’s observed BOLD timeseries, d) HRF-convolved psychological terms reflecting the expected BOLD response to the task of interest as well as all other modeled task events.

For example, again for encoding in the Adult2 cohort, the formula was:

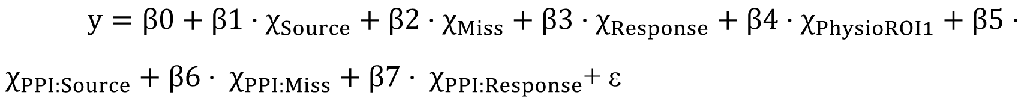

Here, in PPI terms, y is a physiological variable, namely the BOLD time series from the test region (ROI2). The independent variables in this model included one constant term, one regressor of the seed ROI time series (ROI1), three regressors of task activations (Source, Miss, Response), and three corresponding PPI regressors/terms (see **SFig 2** for a matrix representation of the same example, and **STable 1** for specific for across cohorts and memory processes). The β5 estimate was used for the source memory condition. For each subject, the PPI models were built for each ROI and fitted to all other ROIs.

**Figure 2.**
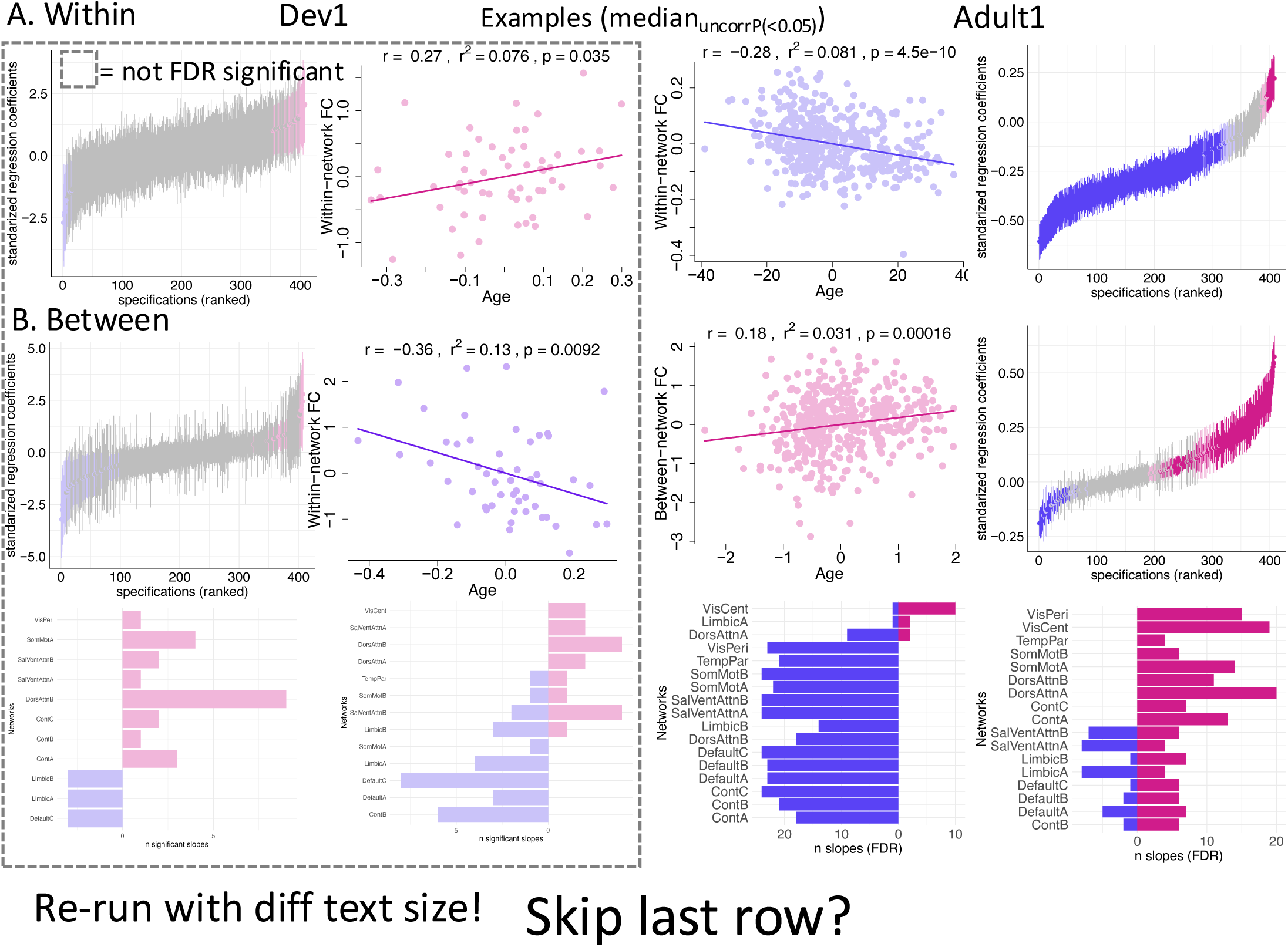
Task-general FC and age. (**A**) Within- and between-network (**B**) task-general FC associations with age for Dev1 and Adult1. Specification curves for associations with age (first and fourth columns). Example scatter plots (median regression coefficient with uncorrected P values < 0.05, FC and age residualized by model covariates for visualization, statistical values in the header reflects associations between these two variables and might differ slightly from the full model). Please note that this median often does not represent the median of all relationships, which instead could be non-significant. (**C**) Involved networks for within- and between-FC, respectively, for Dev1, and (**D**) Adult1.

Following estimation, the regression coefficient associated with a) is taken to reflect region B’s influence on ROI’A during the specified task component. Here, the resulting asymmetric 400 x 400 matrix of PPI effects was finally symmetrized (Di et al., 2017) to allow direct comparison with the other bidirectional FC measures.

The cPPI and gPPI calculation was done using MATLAB 2017a, SPM 12.0, and gPPI toolbox 13.1. For all FC matrices, the resulting correlation coefficients were Fisher-transformed to z-values. We have previously looked at FC between single regions in overlapping samples, both cortico-cortical (Capogna et al., 2022), hippocampal-subcortical (Ness et al., 2022) and hippocampal-cortical connections (Raud et al., 2023), but here we focused on cortical within- and between network FC.

For TSFC, TRFC, cPPI FC, to obtain within- and between-network FC values, we first used the 17-network parcellation (Yeo et al., 2011) to which each region of the 400 Schaefer atlas is assigned (**Fig. 1**). For each network, we obtain one measure of within-network FC, which is the average FC across regions within that network, and one measure of between-network FC, which is the average FC between across the region in the network with all other regions. Hence, for each FC method at encoding and retrieval, respectively, we obtain 17 within- and 17 between-network measures. The networks, with abbreviations, are: Visual A (VisCent), Visual B (VisPeri), Somatomotor A (SomMotA), Somatomotor B (SomMotB), Temporal Parietal (TempPar), Dorsal Attention A (DorsAttnA), Dorsal Attention B (DorsAttnB), Salience/Ventral Attention A (SalVentA), Salience/Ventral Attention B (SalVentA), Control A (ContA), Control B (ContB), Control C (ContC), Default A (DMNa), Default B (DMNb), Default C (DMNc), Limbic A (LimbicA), and Limbic B (LimbicB). In FC analysis, negative/anticorrelated edges can be included, but such edges pose challenges of interpretation. Hence, like previous influential work (Chan et al., 2014), we ran separate analyses, once including negative z-values, and once excluding negative z-values (**Table 2**).

As we used predefined networks, based on rs-fMRI (Yeo et al., 2011), we assessed the presence of the defining features of a network, namely that within-network connectivity was higher than between-network connectivity (Newman and Girvan, 2004). In line with the reasoning that TGFC aligned more closely with rs-fMRI, when we used TGFC, within-network connectivity was higher than between-network connectivity across processing steps and networks in all samples, crucially also for the developmental subsamples (**SFig. 3-4**, P < 0.0001, FDR corrected across networks and memory process (encoding and retrieval), paired t-tests [two-sided, Student’s t-Test if Shapiro-Wilk Normality Test was non-significant, and Mann-Whitney’ test if significant], except LimbicB for Adult2 in 9 (2%) instances. In contrast, predefined network for MMFC (gPPI), did not show higher within-network FC compared with between-network FC for most networks across the four processing steps (**SFig. 5-6**). Instead, between-network FC was higher for most networks, at encoding and particularly at retrieval when investigated using gPPI (P < 0.05, FDR corrected across networks, and encoding and retrieval combined). For MMFC (gPPI), to account for this interesting observation that merits exploration in future work, we therefore used data-driven, modularity-based networks.

**Figure 3.**
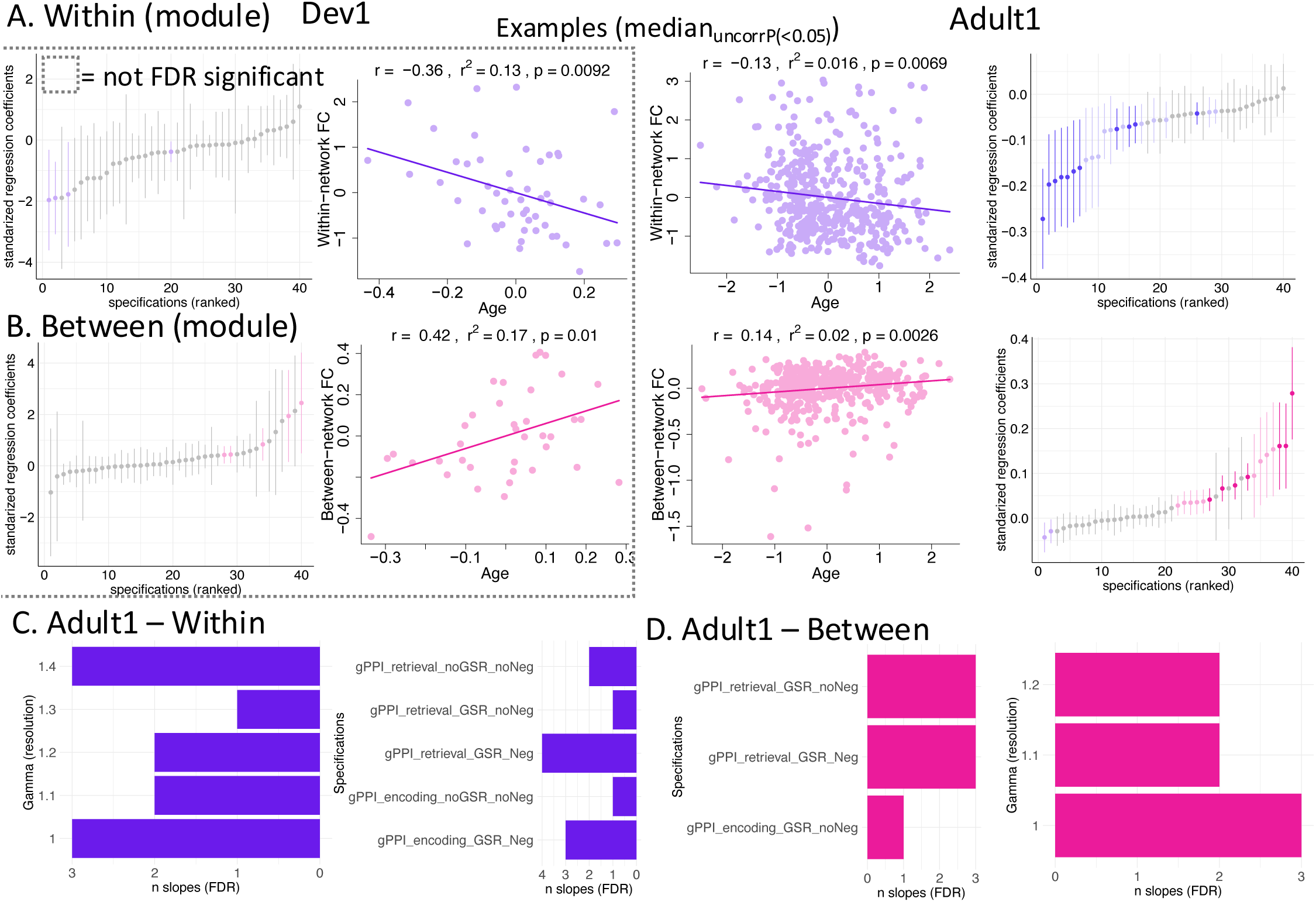
Memory-modulated FC and age. (**A**) within- and (**B**) between-network memory-modulated FC (MMFC) associations with age for Dev1 and Adult1. Specification curves for associations with age (first and fourth columns). Example scatter plots (median regression coefficient with uncorrected P values < 0.05, FC and age residualized by model covariates for visualization). (**C-D**) For Adult1, involved network resolution parameters γ (lower yields fewer networks) and analytic approaches for within- and between-FC, respectively.

**Figure 4.**
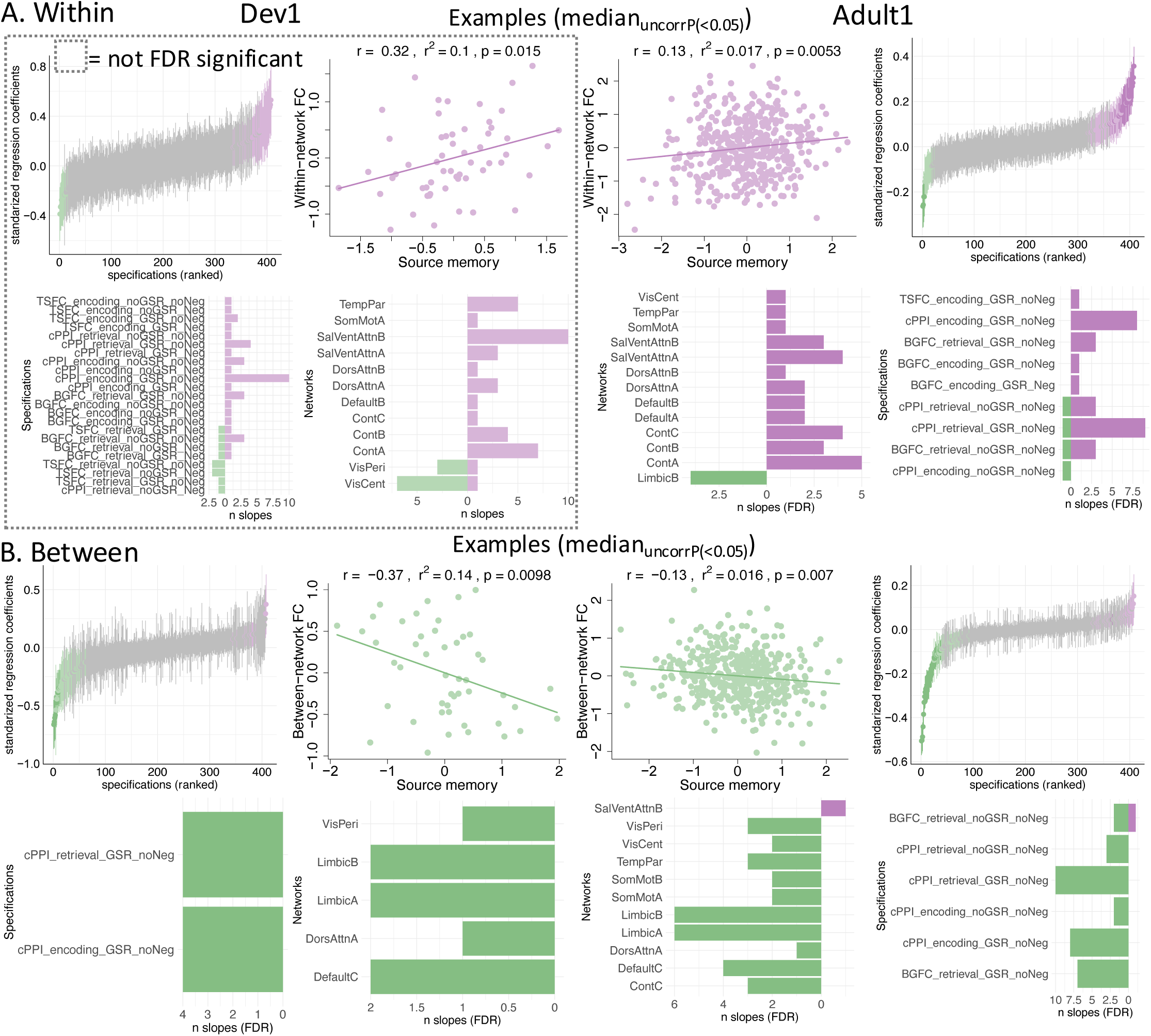
Task-general FC and memory performance. (**A**) Within- and (**B**) between-network task-general FC (TGFC) associations with memory for Dev1 and Adult1. Specification curves for associations with age (first and third row, first and fourth columns). Example scatter plots (first and third row, second and third columns, median regression coefficient with uncorrected P values < 0.05, FC and age residualized by model covariates for visualization). Involved analytic approaches (second and fourth row, first and fourth columns) and networks (second and fourth row, second and third columns). TSFC=Task-state FC. BGFC=Background FC.

**Figure 5.**
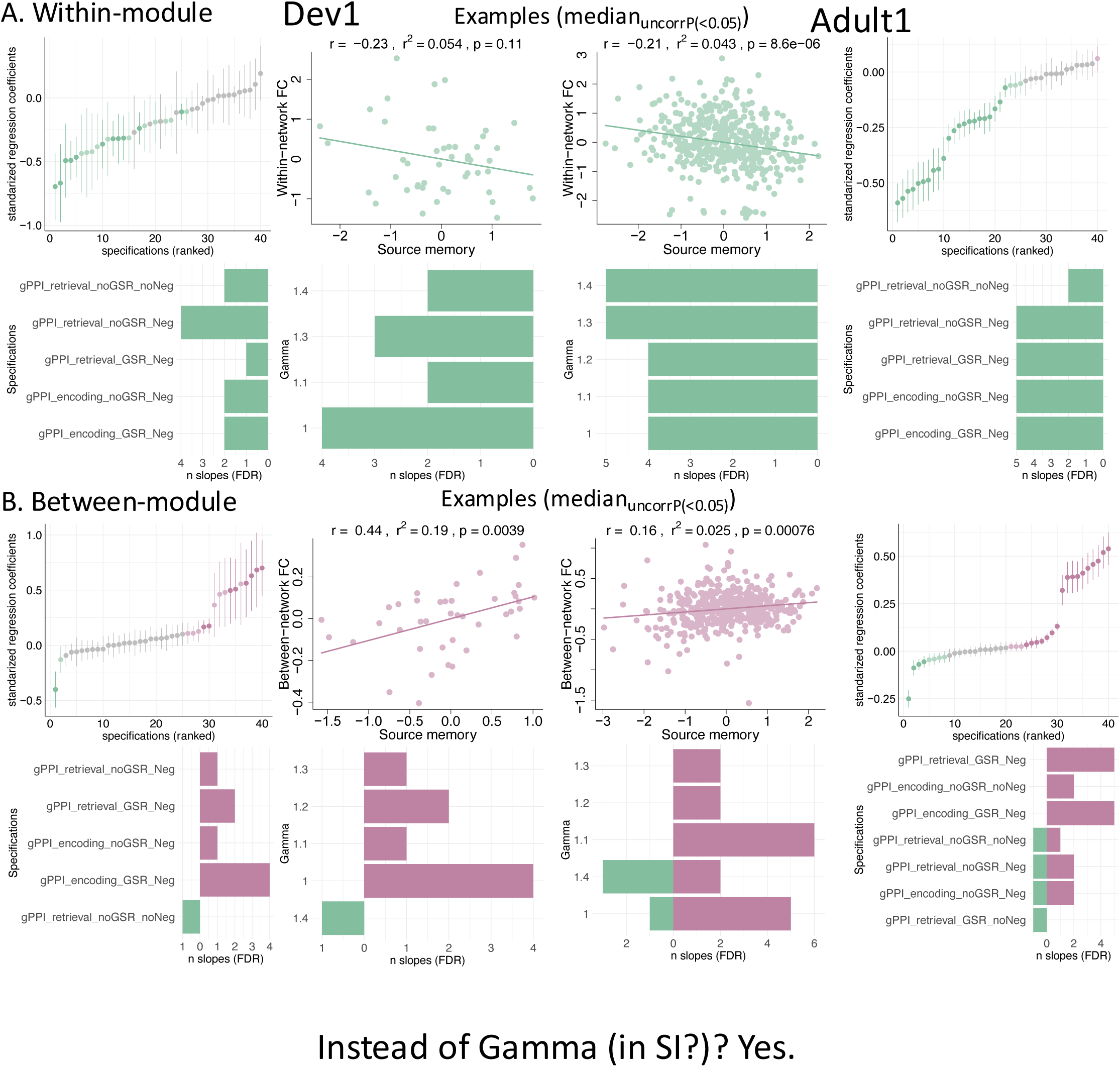
Memory-modulated FC in modularity-derived networks. (**A**) Within- and (**B**) between-network task-general FC (MMFC) associations with memory for Dev1 and Adult1. Specification curves for associations with age (first and third row, first and fourth columns). Example scatter plots (first and third row, second and third columns, median regression coefficient with uncorrected P values < 0.05, FC and age residualized by model covariates for visualization). Involved analytic approaches (second and fourth row, first and fourth columns) and network resolution parameters γ (lower yields fewer networks, second and fourth row, second and third columns).

Data-driven, modularity-based networks were derived using the Louvain network/community detection algorithm (Blondel et al., 2008) using the Brain Connectivity Toolbox (Rubinov and Sporns, 2010) for Python (https://github.com/aestrivex/bctpy). The resolution parameter (γ), was set at the default 1, as well as 1.1, 1.2, 1.3, and 1.4 to detect a number of modules more comparable to the size of the predefined networks. For each participant, we ran the algorithm, at each γ value, across 500 iterations, and retained the community/network corresponding to the solution with highest statistic, and subsequently identified and removed any singleton modules. When excluding negative values, the *louvfunc* term was ‘modularity’, while with negative values included we used ‘negative_asym’ (Rubinov and Sporns, 2011). Due to a varying number of networks per person, values per person were derived as an average across networks (and not on a per-network-basis as for the predefined networks), and number of networks was used as a covariate in these analyses.

The underlying assumption of the importance of cortical within- and between network FC is evaluated by also testing associations from a more specific level of brain resolution, namely (i) FC within the parahippocampal cortices (PHC) and (ii) between the PHC and the rest of the cortex, as PHC are pivotal cortical regions for memory encoding and retrieval (Maass et al., 2018). As we, in line with previous work discussed above, focus on cortical networks we focused on the cortical aspects of the medial temporal lobe (the parahippocampal cortex) instead of subcortical regions (the hippocampus). The regions used were the 5 PHC regions in the Schaefer 400 parcellation PHC_1-3 in the left hemisphere, and PHC_1-2 in the right, all part of the DefaultC network.

### 2.5 Multiverse-inspired analysis and specification curves

Inspired by Bloom et al. (2022), to address the hypotheses, we conducted a multiverse analysis and constructed specification curves. A multiverse analysis involves performing the analysis of interest across the whole set of reasonable scenarios, or so-called specifications. Here, the analyses were linear regression models. For non-predefined networks, we used Cook’s distance, with the cut-off of 1, after visual inspection showing potentially deviant data points. The result displays the stability or robustness of a finding (Steegen et al., 2016). As such, a multiverse analysis can help to show how conclusions vary due to analytic choices.

Analyzing fMRI data entails a host of choices between different approaches. Performing analyses “across the whole set of reasonable scenarios” is impractical, and we took the approach of sampling several reasonable and commonly used analysis choices, as well as across all networks (**Table 2**). Despite being far from comprehensive, and hence potentially better described as a multiverse-inspired analysis, this approach enables a thorough investigation into the stability or robustness the results after our chosen preprocessing methods. Specifically, each analysis consisted of running a model for (i) each of the 4 FC methods, with or without GSR, and with or without excluding negative values, across each predefined network (also repeated in the data-driven networks), for encoding and retrieval (except for Dev2). On a per analysis, per network basis, we defined and excluded outliers as values that was either 1.5 × the interquartile range (IQR) lower than the first quartile, or a value that was 1.5 × IQR greater than the third quartile.

We constructed specification curves by ranking models ascendingly by their regression coefficient (beta) estimates for the term of interest (age and memory, respectively) for interpretation and visualization (Bloom et al., 2022; Simonsohn et al., 2020). We did not conduct formal null hypothesis testing of specification curves. Instead, we reported the proportion of (FDR-corrected) significant specifications. Specification curves plotted with the help of the R package *specr* (Masur and Scharkow, 2020). Although potentially too conservative, as the specifications are related, sometimes quite highly, we also corrected all betas in each specification curve for multiple comparisons using false discovery rate (FDR) via the *p.adjust* function (*stats* package) in R, with the “BY” method (Benjamini and Yekutieli, 2001).

To test whether relationships between FC and memory performance varied with age, we tested for differences between the FC-memory regression coefficients, as presented in the results, obtained separately for Dev1/2 and Adult1/2 (Cohen et al., 2013). This approach enabled comparisons of the actual regression coefficients presented in the results, which was deemed more parsimonious than introducing new models with interaction terms. Similar tests were performed for the FC-age relationships. Specifically, for each test, we calculated the differences between the regression coefficients from Dev1/2 and Adult1/2, respectively. The standard error (SE) of this difference (between the two coefficients) was calculated as:

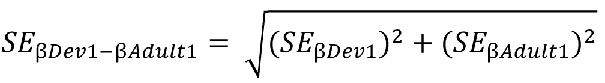

This SE was then used to calculate a Ζ value (as the ratio of the difference between reg coefficients and SE):

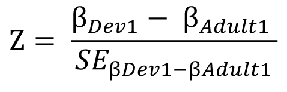

The p values for the Z values were obtained using the *pnorm* function in R.

### 2.6 Out-of-sample FC prediction of memory performance

Inspired by connectivity-based predictive modeling (Shen et al., 2017), we tested, for significant relations between network FC and memory, the predictive, out-of-sample power of network FC on memory via a leave-one-out cross-validation (LOO-CV) approach. Due to (i) the high number of analyses in our multiverse-inspired approach, and (ii) the main focus of the present work was assessing the relationship between FC and memory, we did not perform LOO for FC-age relationships. Specifically, looping over the whole sample, leaving one participant out each time, we re-ran the model on the sample minus the left-out participant (training). From this training sample, we used the resulting model parameters to obtain a predicted memory value for the left-out participant (test). After obtaining predicted values for each left-out participant, we correlated the predicted values with the original values. To test for significance, we calculated a P value by permutation analysis, repeating the analysis 1000 times with randomly rearranged values to obtain a null distribution. The resulting P value was the number of times the actual correlation value (between predicted and original values) was higher than the absolute value of the permuted correlation values. To control for covariates like mean FC, motion, and age, we applied the full model including these covariates for the training, but only used the memory term (and the intercept) to calculate the predicted values.

### 2.7 Data and code accessibility

Data and analysis scripts will be shared at the dedicated project at Open Science Framework. The MRI data for most participants may be available upon reasonable request, given appropriate ethical, data protection, and data-sharing agreements. Requests for the raw MRI data can be submitted to the principal investigator (Prof. Anders Fjell, University of Oslo).

Individual-level data availability may be restricted as participants have not consented to publicly share their data.

## 3. Results

### 3.1 Memory performance

In **Fig. 1D**, we present memory performance as a function of age per sample. As expected, and shown previously Dev1 and Adult1 (Vidal-Pineiro et al., 2021), memory performance was higher with higher age in development (approximately from 10 to 20 years, Dev1: r =.37, P=0.008, Dev2: r=0.16, P=0.06, median r and P values shown as number of participants included differed depending on outliers in a network-specific manner), and lower with higher age in adults (r=-.51 and -.38 for Adult1 and Adult2, respectively, uncorrected P < 0.001).

### 3.2 FC and age

#### 3.2.1 Task-general FC and age

First, we tested associations between age and within- and between-network task-general FC (TGFC), respectively, including covariates of motion, sex, and mean cortical FC. **Table 3** summarizes the percent of significant associations. As shown in **Fig. 2**, for Adult1, we observed, as expected, mainly negative age slopes for within-network FC, and mainly positive slopes for between-network FC. The Dev1 associations were much fewer, and although a slight majority of associations were in the expected, opposite direction, no associations survived FDR correction (**Fig. 2**). Testing the differences in age regression slopes between Dev1 and Adult1, no differences survived FDR correction. The results for Adult2 were similar to Adult1 (**SFig. 7**, the results for Adult2 and Dev2, the sample with less data, are presented in the Supplementary Material due to space constraints.). Dev2 showed negative associations for within-network FC, in contrast to Dev1, while no between-network associations survived FDR correction, similar to Dev1 (**SFig. 7**). Testing the differences in age regression slopes between Dev2 and Adult2, several differences survived FDR correction, namely 85 (42%) within-FC differences, where all Adult2 age slopes were significant, while the Dev2 age slopes were not significant in all but three instances, and 27 (13%) between-FC, where all Adult2 slopes were significant, while no Dev2 slopes were significant.

**Table 3.** Summary of within- and between-network FC associations with age and memory: (predefined) 17 networks and modules.

|  |  | Age |  |  |  | Memory performance |  |  |  |
| --- | --- | --- | --- | --- | --- | --- | --- | --- | --- |
|  |  | Within |  | Between |  | Within |  | Between |  |
|  |  | Pos | Neg | Pos | Neg | Pos | Neg | Pos | Neg |
| **Task-general FC (17-networks)** | Dev1 | noFDR |  | noFDR |  | noFDR* |  | 0 | 2 |
|  | Adult1 | 3 | 77 | 38 | 8** | 7 | 1 | 0 | 8 |
|  | Dev2 | 0 | 9 | noFDR |  | noFDR |  | 0 | 1 |
|  | Adult2 | 1 | 65 | 23 | 4 | noFDR |  | NS |  |
| **Memory-modulated FC (modules)** | Dev1 | noFDR |  | noFDR |  | 0 | 28 | 20 | 3 |
|  | Adult1 | 0 | 28 | 18 | 0 | 0 | 55 | 43 | 10 |
|  | Dev2 | NS |  | noFDR |  | 0 | 50 | 35 | 10 |
|  | Adult2 | noFDR |  | noFDR |  | 0 | 2.5* | 3 | 5 |

Values in percent (%) of tests in the corresponding analysis. Pos/Neg=Positive/Negative associations. noFDR=No relations not surviving FDR correction. NS=no significant associations. \* <1% significant interaction terms within subsample.

#### 3.2.2 Memory-modulated FC and age

We then tested how MMFC related with age (**Table 3**), using the data-driven, modularity- derived networks, or modules, including covariates of motion, sex, mean cortical FC, and number of modules. As for TGFC above, Adult1 showed FDR-significant relations with age, negative for within-FC and positive for between-FC. For Dev1, the few significant associations (10 and 13%) did not survive FDR correction (**Fig. 3**). Testing the differences in age regression slopes between the Dev1 and Adult1, no associations survived FDR correction. Dev2 and Adult2 showed 6 significant associations in total, none surviving FDR correction (**SFig. 8**), with no FDR-significant differences between age regression slopes.

#### 3.2.3 Parahippocampal cortex (PHC) FC and age

For TGFC from the PHC, three samples showed associations with age (**Table 4**). For Adult1, Adult2 and Dev2, associations were strongest and most consistent for within-PHC connectivity (100%, all negative), but also present for between-PHC, that is, the mean of the mean FC between each PHC region and the rest of the cortex. For MMFC, we observed only 3 associations in the Adult1 subsample.

**Table 4.** Summary of PHC associations with age and memory (within-PHC and between PHC and rest of cortex).

|  |  | Age |  |  |  | Memory performance |  |  |  |
| --- | --- | --- | --- | --- | --- | --- | --- | --- | --- |
|  |  | Within |  | Between |  | Within |  | Between |  |
|  |  | Pos | Neg | Pos | Neg | Pos | Neg | Pos | Neg |
| **Task-general FC** | Dev1 | noFDR |  | noFDR |  | NS |  | 0 | 4 |
|  | Adult1 | 0 | 100 | 17 | 13 | noFDR |  | 0 | 17 |
|  | Adult2 | 0 | 100 | 0 | 17 | noFDR |  | noFDR |  |
|  | Dev2 | 0 | 100 | 0 | 17 | NS |  | 0 | 8 |
| **Memory-modulated FC** | Dev1 | noFDR |  | NS |  | noFDR |  | NS |  |
|  | Adult1 | 13 | 0 | 13 | 13 | NS |  |  | 13 |
|  | Dev2 | NS |  | NS |  | NS |  | noFDR |  |
|  | Adult2 | NS |  | noFDR |  | 50 | 0 | 25 |  |

Values in percent (%) of tests in the corresponding analysis. Pos/Neg=Positive/Negative associations. noFDR=No relations not surviving FDR correction. NS=no significant associations.

### 3.3 FC and memory performance

#### 3.3.1 Task-general FC and memory performance

For within-network TGFC and memory performance (**Table 3**), Adult1 showed 33 associations, mainly positive (**Fig. 4A**) where within-network FC were lower with higher memory performance. In all tests, we included the covariates of age, sex, motion, and mean cortical FC. None of these relations yielded significant, LOO-CV predictions (maximum r=.41, P_perm_=0.075). Also, we observed 3 interactions with age, where only the younger, and not older adults, showed a positive relation (the association in the older was not significant when tested separately, groups based on median age split). For Dev1 no relationships survived FDR correction (**Fig. 4A**), but six of the non-corrected relationships showed LOO- CV predictions: 3 in ContA, 2 in ContB, and in SalVentAttnB (**SFig. 9**).

For between-network TGFC and memory performance, Dev1 and Adult1 showed a total of 41 associations, mostly negative (**Table 3 and Fig. 4B**). For Adult1, 10 LOO-CV predictions were significant (**SFig. 9**), while for Dev1, twenty-two associations (also non-FDR-significant ones) showed LOO-CV predictions. Testing the differences between Dev1 and Adult1 memory regression slopes, no associations for within- or between-FC survived FDR correction.

For Dev2 and Adult2 combined, 1 FDR-significant association was observed (**SFig. 10**), but the prediction was not significant (r=.57, P_perm_= 0.11). Testing the differences between Dev2 and Adult2 memory regression slopes, no differences survived FDR correction.

#### 3.3.2 Memory-modulated FC and memory performance

Using the average within- and between-network FC of data-driven, modularity-derived networks, both Dev1 and Adult1 showed only negative within-MMFC associations with memory (**Table 3** and **Fig. 5A**), across all levels of resolution tested (γ = 1-1.4). LOO-CV tests for Adult1 (**SFig. 11A**) showed five significant predictions. For Dev1, we observed five significant LOO-CV predictions. For between-network MMFC, both Dev1 and Adult1 showed mainly positive associations (**Fig. 5B**). LOO-CV tests for Adult1 (**SFig. 11B**), showed 2 negative and 5 positive associations were significant predictions. For Dev1, FC from 1 negative and 5 positive associations also yielded significant LOO-CV predictions.

Testing the differences in memory regression slopes between Dev1 and Adult1, no associations for either within- or between-FC survived FDR correction.

The findings were mainly replicated in Dev2 and Adult2 (**Table 3** and **SFig. 12**), except for Adult2 showing only 3 associations for between-FC and memory, with 2 being negative. LOO-CV tests (**SFig. 13**) showed no significant predictions for Adult2, while for Dev2, four within-FC predictions, and two between-FC predictions, were significant. Testing the differences in memory regression slopes between the Dev2 and Adult2 for the encoding phase, 1 (5%) within-FC survived FDR correction with the Dev2 associations being negative and FDR-significant, while the Adult2 was positive, but not significant, and 1 (5%) between- FC (both associations positive, but only FDR-significant in Adults2).

#### 3.3.3 PHC FC and memory

In the parahippocampal cortices (**Table 4**), no subsample showed associations with memory for within-TGFC. For between-TGFC, a total of 6 associations were observed in Adult1, Dev1 and Dev2. Specifically, for Adult1, we found 4 negative associations, with 1 LOO-CV predictions for retrieval (r=. 0.70, P_perm_ = 0.001). Both Dev1 and Dev2 showed 1 negative association each, but no prediction (r=.47, P_perm_=0.098 and r=.44, P_perm_=0.175, respectively). For between-MMFC and memory, Adult1 showed 1 negative association for retrieval, with a LOO-CV prediction of memory scores: r=.53, P_perm_=0.031. Adult2 showed 4 positive association for within-MMFC and 1 positive a positive association for between-MMFC, but with no predictions (P_perm_>0.40).

## 4. Discussion

Investigating network functional connectivity (FC) across the lifespan during both encoding and retrieval of episodic memories, we found (1) that memory-modulated FC showed relations with memory performance in both development and aging, including the ability to predict out-of-sample memory scores. In 97.5% of the cases, these memory associations were age-invariant, implying that regardless of their age, persons with low network differentiation showed worse cognitive performance than similarly aged individuals with higher levels of differentiation (Koen and Rugg, 2019). (2) We also found that FC reflecting more condition-unspecific, task-general properties (task-general FC) showed (i) lower segregation with higher age in adults, and (ii) associations with memory performance in an age-invariant fashion. For task-general FC in the largest adult sample, lower segregation related to worse memory, in line with the dedifferentiation view. However, in development, as both task-general and memory-modulated FC showed only weak signs higher segregation with higher age, we observed few signs of differentiation, and hence at least partly a lack of support for a differentiation-dedifferentiation lifespan pattern (Tucker-Drob, 2009). If these results are replicated and extended using longitudinal studies (Chong et al., 2019), theories of development and aging need to incorporate why lower network segregation in aging, accompanied by lower memory, do not have an opposite counterpart in development, that is, gradually higher segregation with development accompanied by higher memory performance. As relationships were tested in a multiverse-inspired approach across different analytic choices using specifications curves, it is noteworthy that most tests showed few and weak associations, and that several significant results differed based on analytic choices. As such, the results highlight that analytic choices may influence the inferences drawn in crucial ways. Overall, the main result, based mainly on the cohort with most data, was similar relations between FC and episodic memory performance from age 7 to 82 years. As age-invariant relations might indicate fundamental aspects of brain organization (Rugg, 2017), this result suggest that network segregation might be pivotal for memory accuracy across much of the healthy lifespan.

### 4.1 Network segregation and memory: similar relations from 7 to 82 years

The associations between FC and memory performance across all analyses was mainly in the same direction for adults and development, pointing to age-invariant relations between network segregation and episodic memory across 7 to 82 years of age. This means that the sign and strength of the relations show little variation with age (Koen et al., 2020). As discussed by Srokova et al. in relation to neural selectivity (Srokova et al., 2020), although an age-invariant relation does not rule out a role for segregation in mediating age-related cognitive decline, it does suggest that the contribution of within- and between-network connectivity to memory performance is stable. As argued by Rugg (2017), the finding of age-invariant relations between a cognitive process and a pattern of brain activity can support the notion that the relationship represents a fundamental aspect of functional brain organization. Hence, network segregation might be an important determinant of memory accuracy both in development and aging (Rugg, 2017).

In development, we observed relationships between memory and both within- and between-network MMFC. Although Petrican et al. (2017) reported association between within-network FC and working memory performance in development (12-21 years), they did not find associations for episodic memory. In adults, the findings of memory relations are in line with several rs-fMRI studies (Chan et al., 2014; Stumme et al., 2020; Varangis et al., 2019), linking FC with memory outside scanner. Specifically, in Varangis et al. (2019), the link was with mean FC and no network showed consistent relations with memory performance. Relationships were not tested on a per network basis in Chan et al. (2014), while the well-powered study by Stumme et al. (2020) found both within- and between-network links with “non-verbal memory and attention”, and a lower between-default and frontoparietal network link with “verbal memory and fluency”. Here, we expand previous findings by showing how segregation metrics also yield out-of-sample predictions of memory performance for both within- and between-network FC in both adults and development.

### 4.2 Network segregation and memory: differences in differentiation across the lifespan?

In adults, task-general FC showed that lower segregation with age (lower within- and higher between-network FC) related to lower memory performance. This pattern is often attributed to dedifferentiation (Chan et al., 2014; Koen and Rugg, 2019). In contrast, one previous study using cPPI based on source memory fMRI data, interpreted the results as beneficial integration reflecting compensation, as more outside-module connections in a medial temporal lobe module associated with better source memory (Monge et al., 2018). They argued that evidence for dedifferentiation would have shown that the higher between-network connection associated with worse source memory performance. Here, we observed this pattern (higher between FC, lower memory) for task-general FC (including cPPI) for instance for DMNc (which includes MTL regions) during both encoding and retrieval. Our MTL-specific, parahippocampal (PHC) analysis showed similar results (albeit less consistently).

For memory-modulated FC, we observed results in accordance with Monge et al. Hence, our findings mainly support a dedifferentiation view, but are partly also in line with a compensation view. However, as we observed mainly similar patterns in development (albeit less strong), the results indicate that age-related differences in FC are not necessarily aging-specific, and hence not a compensation process, but might instead reflect processes with a developmental origin.

### 4.3 Network segregation and age: few signs of higher segregation after 7 years of age

Task-general FC in adults showed less segregation with higher age across the majority of networks and processing choices, as expected from the rs-fMRI literature (Chan et al., 2014; Setton et al., 2022; Stumme et al., 2020; Varangis et al., 2019; Zonneveld et al., 2019) and background task FC (Grady et al., 2016; Monge et al., 2018; Rieck et al., 2021). In development, we found similar signs of lower within-network FC with age in one sample (age 10-16 years), which partly echoes Betzel et al. (2014). In a lifespan sample (aged 7-85 years), they found linear negative relations with age in most within-network connections, and positive between-networks relations, that is, a lower segregation with higher age. Here, we did not see indications of higher segregation with higher age in development for any measure, including PHC FC. In development, the predefined networks also showed the defining network properties of higher within-network FC and lower between-network FC per network. This result is therefore likely not explained by lower fit of the predefined networks, defined in adults, in the developmental samples. The finding raises the question of when in development higher segregation starts dominating. Bruchhage et al. (2020), using seed analyses in infants and children (aged 3 months to 6 years), observed that rs-fMRI FC within the same networks was higher at higher ages, and that FC between different networks was lower. This pattern fit higher within- and lower between-network FC with higher age. Hence, like cortical thickness development (Walhovd et al., 2017), segregation might plateau at an early age (< 5 years).

However, like results here, in Bruchhage et al. (2020), there was a fair number of non-significant links as well. A partly alternative explanation is that the within- and between network delineation might be somewhat simplistic, with networks showing more heterogeneous connectivity patterns (Petrican et al., 2017). For instance, Barber et al. (2013) reported that children (8-13 years), compared with adults (20–47 years), had lower within-network FC in circumscribed regions rather than across the whole network. Testing this hypothesis necessitates a sample including children below 5 years and FC measures with a finer resolution.

### 4.4 Network segregation and age: varying associations for memory-modulated FC

The memory-modulated FC did not fit well with the predefined networks. This task-related reconfiguration of networks is in line with previous finding showing important differences between network organization during rest and task states, consistent with the idea that considerable reorganization occurs during specific tasks (Najafi et al., 2016). This fits the notion that a partition of brain regions into nonoverlapping networks is likely an oversimplification of functional brain network organization” (Huang et al., 2024). Instead, task-FC might contain a combination of more stable components, and components that are more state-dependent (Buckner et al., 2013), yielding different, dynamic network configurations. Hence, the use of pre-defined networks for the state-dependent components, as MMFC here, is likely suboptimal.

As a result, we used data-driven networks for memory-modulated FC. Despite this being a somewhat different analysis, memory FC showed similar results as task-general FC, that is, lower within and higher between FC with age. That said, there was a fairly low number of age relations for memory-modulated FC. One possible reason is that most of the age variance was captured by task-general FC. The within-network finding just discussed for memory-modulated FC does not support this alternative, as it showed fairly robust age relations. Another possibility is that the gPPI analyses lacked power, creating false negatives, a common caveat of this type of analysis (O’Reilly et al., 2012). The opposite results warrant caution in the interpretations, and future studies using memory FC (gPPI) would benefit from using data-driven, functional units (Salehi et al., 2020).

### 4.5 Network segregation and memory: direction of associations

We observed both positive and negative FDR-significant relations with memory. Positive relations fit previous studies showing how segregation is lower at higher ages, and that this lower segregation relates to lower memory performance (positive FC memory relation) (Chan et al., 2014). However, studies also find the opposite relations. For instance, Grady et al. (2016) observed that higher between-network FC of the frontoparietal control network was beneficial to memory performance in older but not younger adults. In line with this, Cassady et al. found that lower segregation associated with *better* memory ability in older adults (Cassady et al 2021). Hence, it is somewhat ambiguous in which direction within- and between-network FC are associated to memory at different stages of the lifespan. Likely, the pattern also varies across networks, with sensory-motor networks often showing relations, both with age (Zonneveld et al., 2019) and cognition (Stumme et al., 2020), in the opposite direction compared with association networks, although here we only saw such opposite relations with age.

Another relevant point for interpreting these directional differences, is the use of GSR and the handling of negative values. Although other factors might also contribute, we find different directions in the same networks particularly depending on GSR and whether we include negative values or not. The relations seen with memory were often, although not exclusively, seen when data had been processed using GSR and excluding negative values.

This increased power to detect relations might be due to beneficial effects of GSR in removing global artifacts present in most fMRI scans, which correlate with age and cognitive variables (Power et al., 2017a). Here, we particularly see an effect of GSR in combination with excluding the negative values. As noted by Aquino et al. (2020), GSR alters the distribution of correlations to be centered on zero, and not only by a simple, relative shift of the original distribution. Instead, GSR can alter the correlations in a spatially heterogeneous way, depending on the initial size and strength of correlations between clusters of voxels (Glasser et al., 2018; Saad et al., 2012), although the degree of this effect may depend on the dimensionality of the data Power et al. (2017a). The subsequent removal of the negative values would be a kind of thresholding, a common approach in network science, but due to the shift in distribution of FC values mentioned above, it is not necessarily so here. Thus, these steps could affect the associations seen here only when using GSR and excluding negative values. Alternatively, depending on the dimensionality of the data, we here look at meaningful subsets of the connections, and the associations might be due to beneficial effects of GSR, which also has been reported to improve correlations between FC and behavior (Li et al., 2019). Future analyses would likely benefit from comparing different and more sophisticated denoising approaches, like temporal ICA (Glasser et al., 2018), *rapidtide* (Frederick, 2016-2022) and *DiCER* (Aquino et al., 2020).

### 4.6 Network segregation and memory: mostly non-significant associations

For the memory associations, except for the most highly powered Adult1, there were very few relations, including for mean- and PHC FC. As most relations were seen in the Adult1 sample, which had by far the most participants, some of these null findings may be due to lack of statistical power. Particularly the developmental cohort Dev1 had relatively few participants (<50), which might have resulted in weaker or fewer relationships, or both.

However, we also observed relatively few associations also in the more highly powered Adult2 and Dev2, pointing to additional reasons for the null findings. In a recent study of the same Adult2 and Dev2 cohorts, task-general FC (cPPI) between the hippocampus and cortical regions did also not show associations with memory performance (Raud et al., 2023). One possibility is that the differences in task requirements between Adult1/Dev1 and Adult2/Dev2 influenced results, for instance, the presence of a forced choice task in Adult2 after encoding, could have made the retrieval somewhat less demanding, resulting in less variation and power to detect interindividual differences. Of note, however, since particularly the memory-modulated FC stem from successful encoding and retrieval, the FC assessed here is inherently linked to memory processing despite lack of memory performance associations. That said, although we for Dev2 and Adult2 also observed results in line with age-invariance, the lack of associations for Adult2, makes it challenging to make strong inferences. However, in only two instances (2.5%) did we observe support for age-*variance*, that is, differences between Dev2 and Adult2 FC-memory performance slopes. Overall, although 97.5% of memory association did not show differences between developmental and adult cohorts, the conclusions about memory age-invariance are therefore based on a subset of data, and mainly from the Adult1 and Dev1 cohorts. Further studies could compare associations between different tasks in more similarly powered cohorts.

Further, we did not do inferential analysis on the specification curves, which gives equal weight to all specifications (Simonsohn et al., 2020). While all specifications were justified, statistically valid and non-redundant, we nevertheless are open to some tests being “superior” to others. For instance, for a given sample, tests of a certain kind might be more optimal in making the data closer their “ground truth”, like the use of GSR or not, in different samples with varying degrees of artifacts. Ruling out the link between network FC and memory due to a lack of relationships in other tests would then be ill-suited. Instead, few significant associations then indicate a lack of stability and robustness in the results. Still, interpretations of a minority of significant tests should be done with great caution and tested further in new samples. Second, specification curve analysis merely reduces and does not eliminate ambiguity (Simonsohn et al., 2020). Here, the aim was to present analyses across a host of methods and networks, without selecting *a priori* main and sensitivity analyses. As such, the approach explicitly to avoid selective reporting, but the focus on significant results still run that risk. By showing all results, we believe this approach increases the transparency of the analyses (Steegen et al., 2016), and although not preregistered, avoids selective reporting of results that fit the hypotheses.

### 4.7 Limitations

Dev1 could have been more densely sampled, while Dev2 would have included a somewhat wider age-range. The cross-sectional data does not allow for inferences of change.

Longitudinal FC has shown no relations between change in memory and change in segregation in older adults (n=72, 59–85 years) (Chong et al., 2019), and longitudinal studies are necessary to identify whether associations are driven by between-person differences in FC and memory performance or also by within-person change. The data-driven, modular approach revealed interesting results, but due to the scope of this work, the investigation into further details influencing the modular structure was limited. Although we related gPPI with memory-modulated FC, as noted by Greene et al. (2023), the beginning and end of a task manipulation are unlikely to clearly demarcate the beginning and end of discrete behavioral states. Also, although gPPI might be more sensitive to task-evoked connectivity changes, as supported by the results here, the temporally slower-moving BOLD signal may be less sensitive to faster neural state changes. This is particularly so in memory tasks where the encoding and retrieval phase might evolve over several seconds and are therefore difficult to locate temporally.

## 5. Conclusions

In sum, the main result was the age-invariant associations between FC and memory, suggesting that network relationships may be pivotal for memory accuracy across the healthy lifespan. However, most tests showed few and weak relationships, mainly in the cohort with most data, and several significant results differed based on analytic choices. As such, the results showed how different analytic choices may influence the inferences drawn in crucial ways.

## Supporting information

Supplemental Table [STable] 1

## Acknowledgements

Funding: This work was supported by the Research Council of Norway (grant number #325415).

