## Supplemental Table [STable] 1 for "Network Segregation During Episodic Memory Shows Age-Invariant Relations with Memory Performance From 7 to 82 Years"

### Supplementary Methods

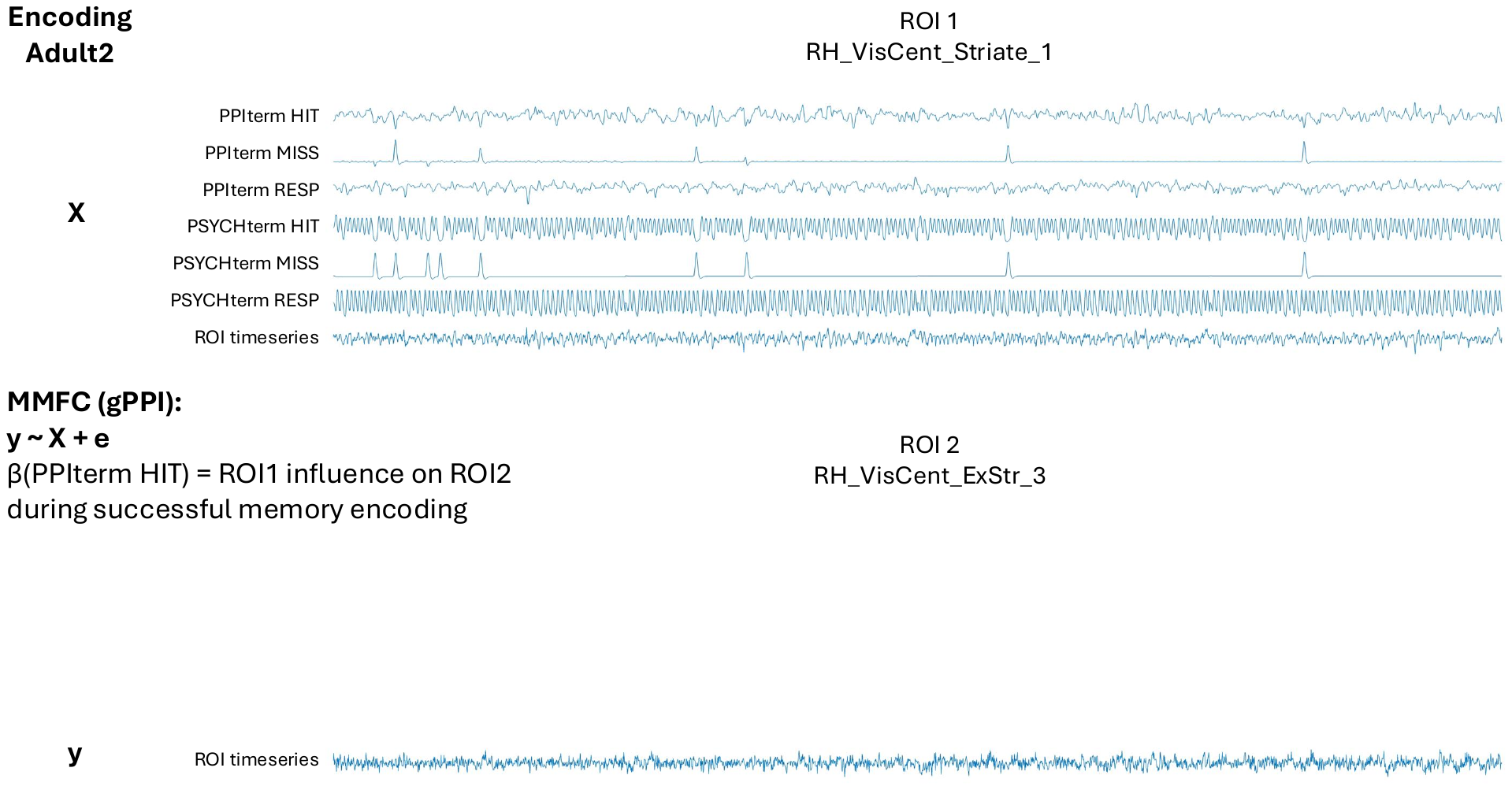

#### Figure S1. Example PPI matrices, gPPI

Example PPI matrices for one participant: Adult2, encoding, gPPI terms.

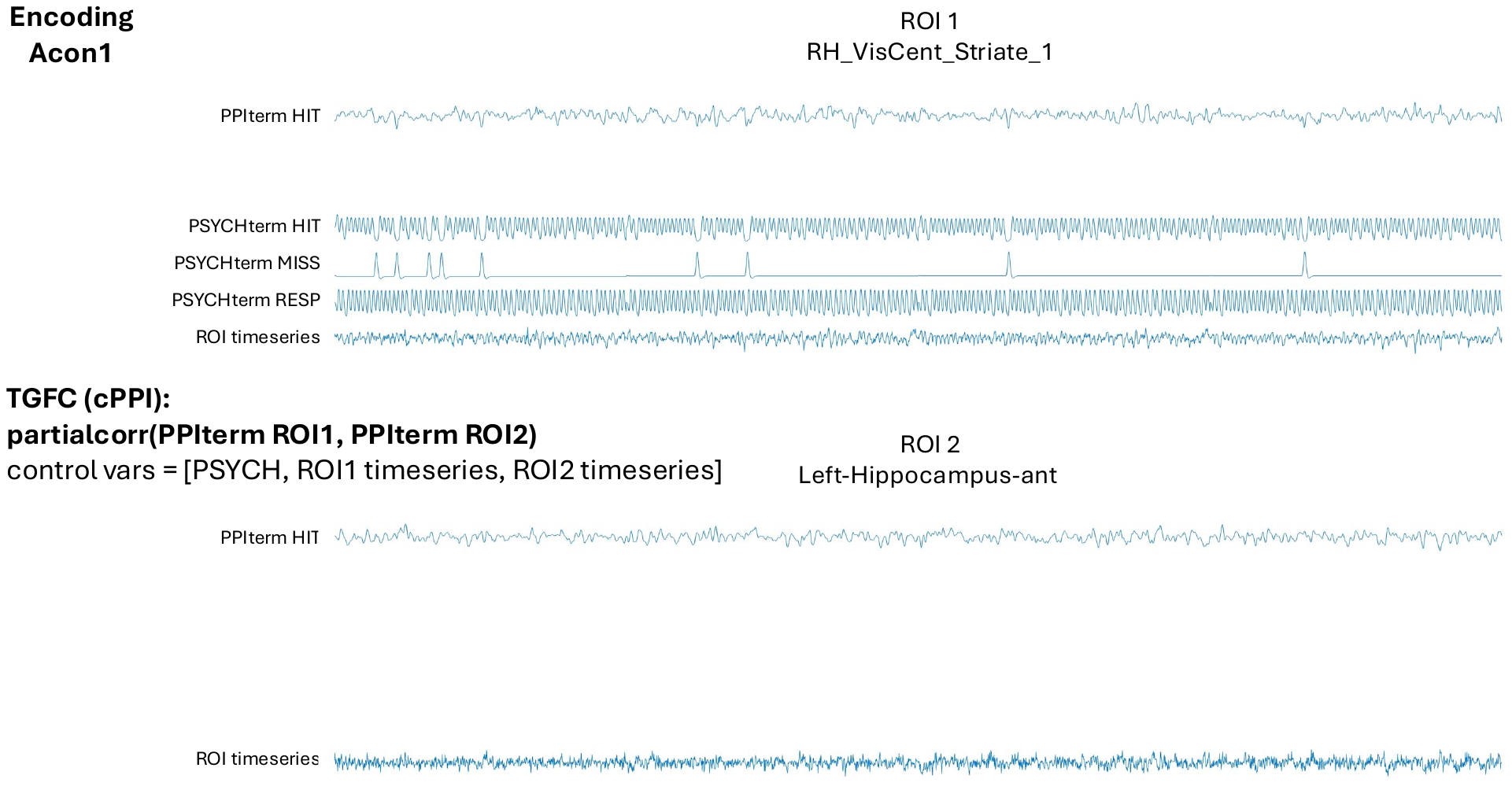

#### Figure S2. Example PPI matrices, cPPI

Example PPI matrices for one participant: Adult2, encoding, cPPI terms.

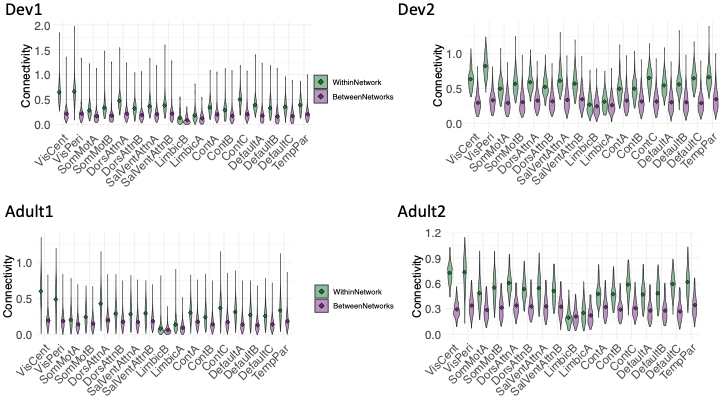

#### Figure S3. Violin plots: task-general FC, encoding, no GSR, negative values

Per network, within- (green) and between-network (violet) FC for task-general FC **(**cPPI) at encoding, no GSR, including negative values.

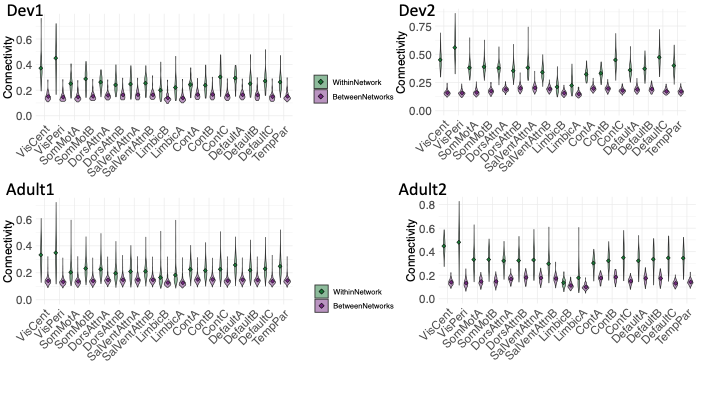

#### Figure S4. Violin plots: task-general FC, encoding, GSR, negative values excluded

Per network, within- (green) and between-network (violet) FC for task-general FC (cPPI) at encoding, GSR, excluding negative values.

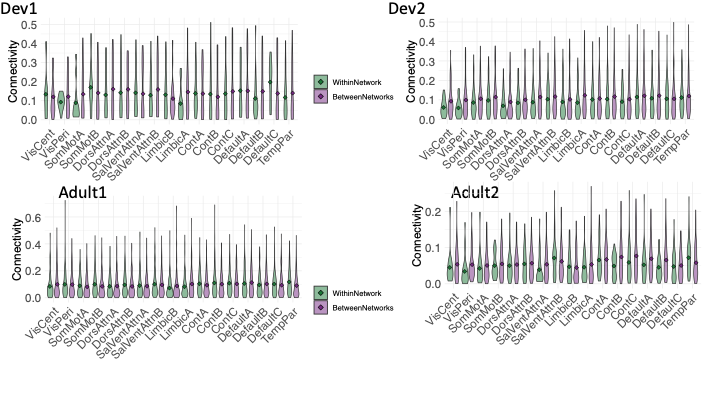

#### Figure S5. Violin plots: memory-modulated FC, encoding, no GSR, including negative values.

Per network, within- (green) and between-network (violet) FC for memory-modulated FC at encoding, no GSR, including negative values.

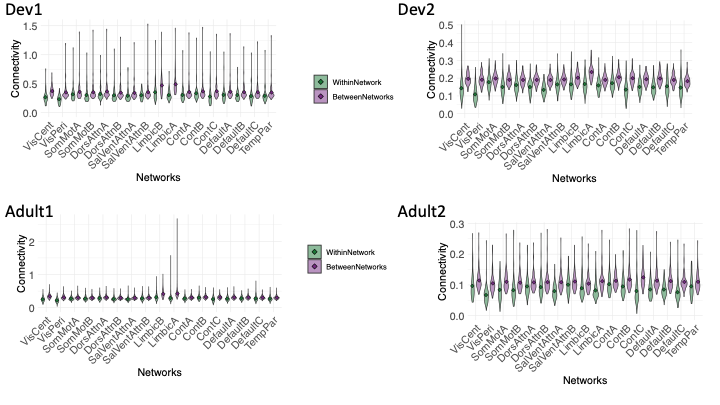

#### Figure S6. Violin plots: memory-modulated FC, encoding, GSR, negative values excluded.

Per network, within- (green) and between-network (violet) FC for memory-modulated FC at encoding, GSR, excluding negative values.

#### Table 1. Model terms for cPPI and gPPI.

|  | **cPPI** | | | **gPPI** | | | |
| --- | --- | --- | --- | --- | --- | --- | --- |
|  | partialcorr(PPI:ROI1,PPI:ROI2, covariates) | | | y ~ X + e | | | |
|  | PPI | Covariates | | y | X |  |  |
|  |  | PSY | PHY | PHY | PPI | PSY | PHY |
| **Dev1, Adult1** | | |  |  |  |  |  |
| Enc | Source | Subseq: Source, Item, Miss. NoResp | ROI1 ROI2 | ROI2 | Subseq: Source, Item, Miss. NoResp | Subseq: Source, Item, Miss. NoResp | ROI1 |
| Ret | Source | Source, Item, Miss, CR, FA, Q2, Q3, Resp | ROI1 ROI2 | ROI2 | Source, Item, Miss, CR, FA, Q2, Q3, NoResp | Source, Item, Miss, CR, FA, Q2, Q3, NoResp | ROI1 |
| **Dev2** | |  |  |  |  |  |  |
| Enc | Hit | Subseq: Hit, Miss. Resp. | ROI1 ROI2 | ROI2 | Subseq: Hit, Miss. NoResp | Subseq: Hit, Miss. NoResp. | ROI1 |
| **Adult2** | |  |  |  |  |  |  |
| Enc | Source | Subseq: Source, Miss. Resp | ROI1 ROI2 | ROI2 | Subseq: Source, Miss. Resp. | Subseq: Source, Miss. Resp. | ROI1 |
| Ret | Source | Source, No Source, New, Resp. | ROI1 ROI2 | ROI2 | Source, No Source, New, Resp. | Source, No Source, New, Resp. | ROI1 |

PPI=the psychophysiological interaction variable (PSY*PHY [deconvolved BOLD timeseries]), PSY= the psychological variable (task design regressor), PHY=the physiological variable (convolved BOLD time series).

Resp=Response. CR=Correct Rejection, FA=False Alarm, Q2=Retrieval question 2 [“Can you remember what you were supposed to do with the item?”], Q3= Retrieval question 3 [“Were you supposed to eat it or lift it?”], NoResp=No response.

### Supplementary *Results*

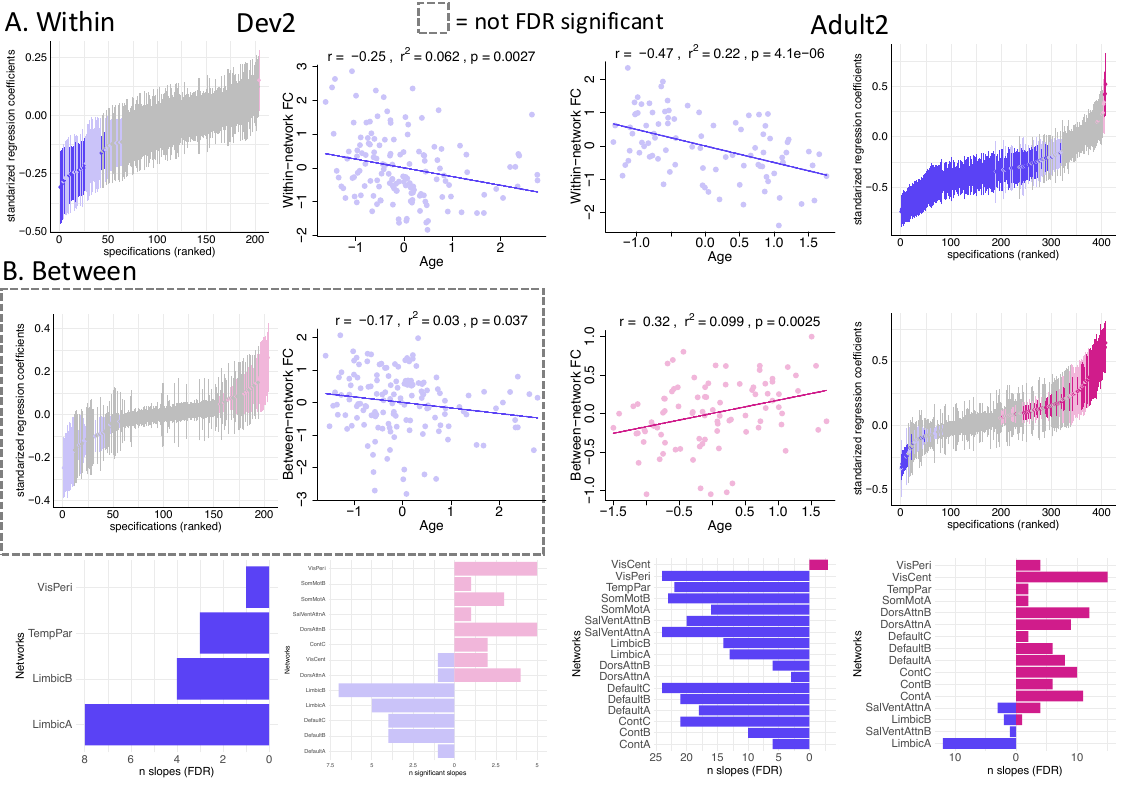

#### Figure 7. Task-general FC and age – Dev2 & Adult2

(**A**) Within- and (**B**) between-network task-general FC for Dev2 and Adult2. Specification curves for associations with age (first and fourth columns). Example scatter plots (median regression coefficient with uncorrected P values < 0.05, FC and age residualized by model covariates for visualization, statistical values in the header reflects associations between these two variables, and might differ slightly from the full model). (**C**) Involved networks for within- and between-FC, respectively, for Dev2, and (**D**) Adult2.

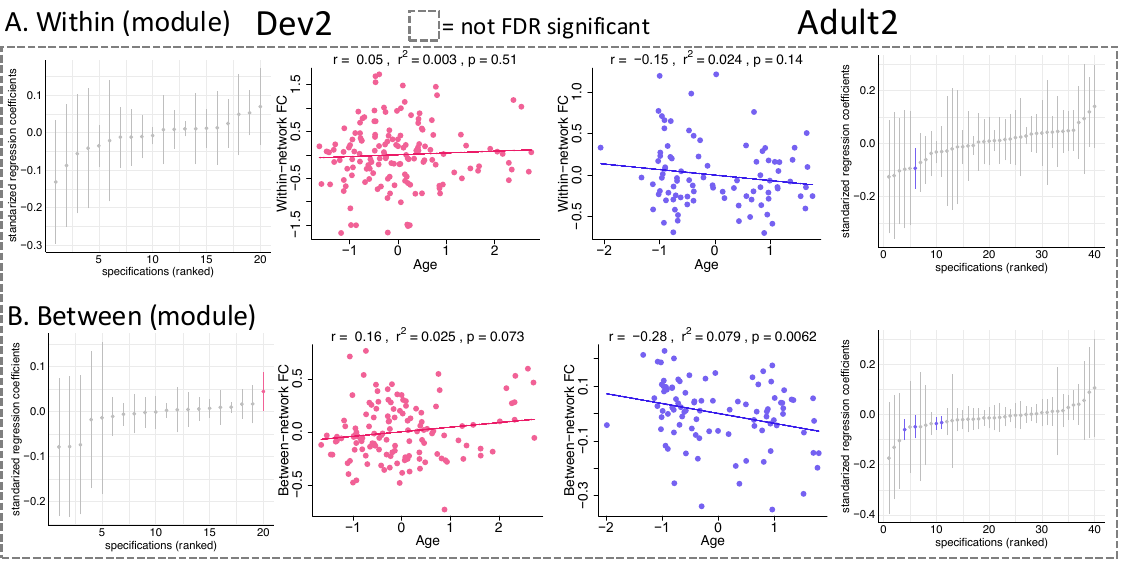

#### Figure 8. Memory-modulated FC and age – Dev2 & Adult2

(**A**) Within- and (**B**) between-network memory-modulated FC for Dev2 and Adult2.. Specification curves for associations with age (first and fourth columns). Example scatter plots of median regression coefficient with uncorrected P values < 0.05 (FC and age residualized by model covariates for visualization).

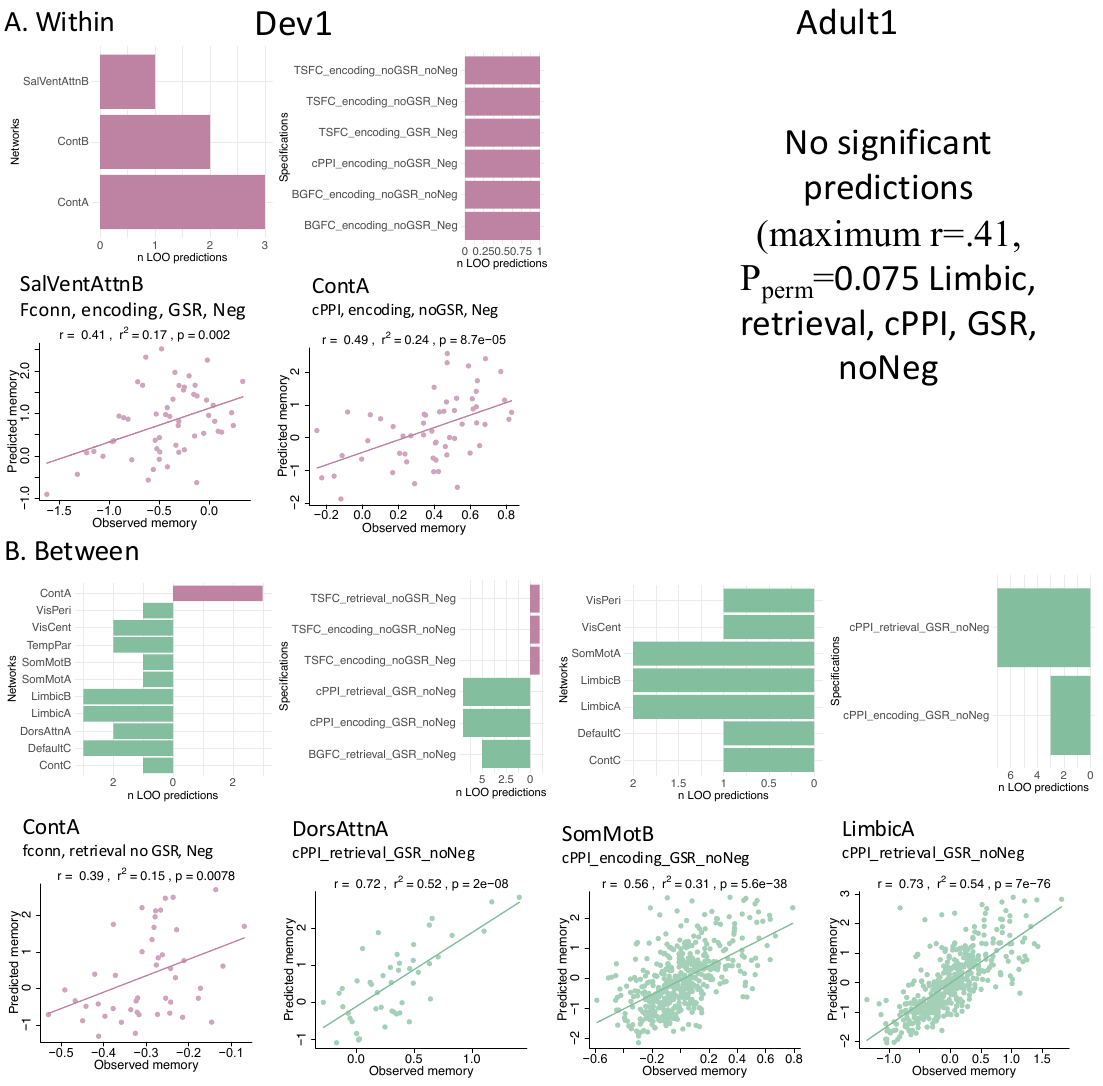

#### Figure 9. Task-general FC and memory *prediction* Dev1 and Adult1

**A**. Within- and (**B)** between-network, task-general FC and memory predictions in Dev1 and Adult1: significant networks (first row, first column; P_perm_ < 0.05), corresponding specifications (first row, second column), and, in the network showing the weakest (first and third column) and strongest (second and fourth column) associations (second and fourth rows), the scatter plot between observed memory performance and predicted memory performance based on FC. TSFC=Task-state FC. BGFC=Background FC.

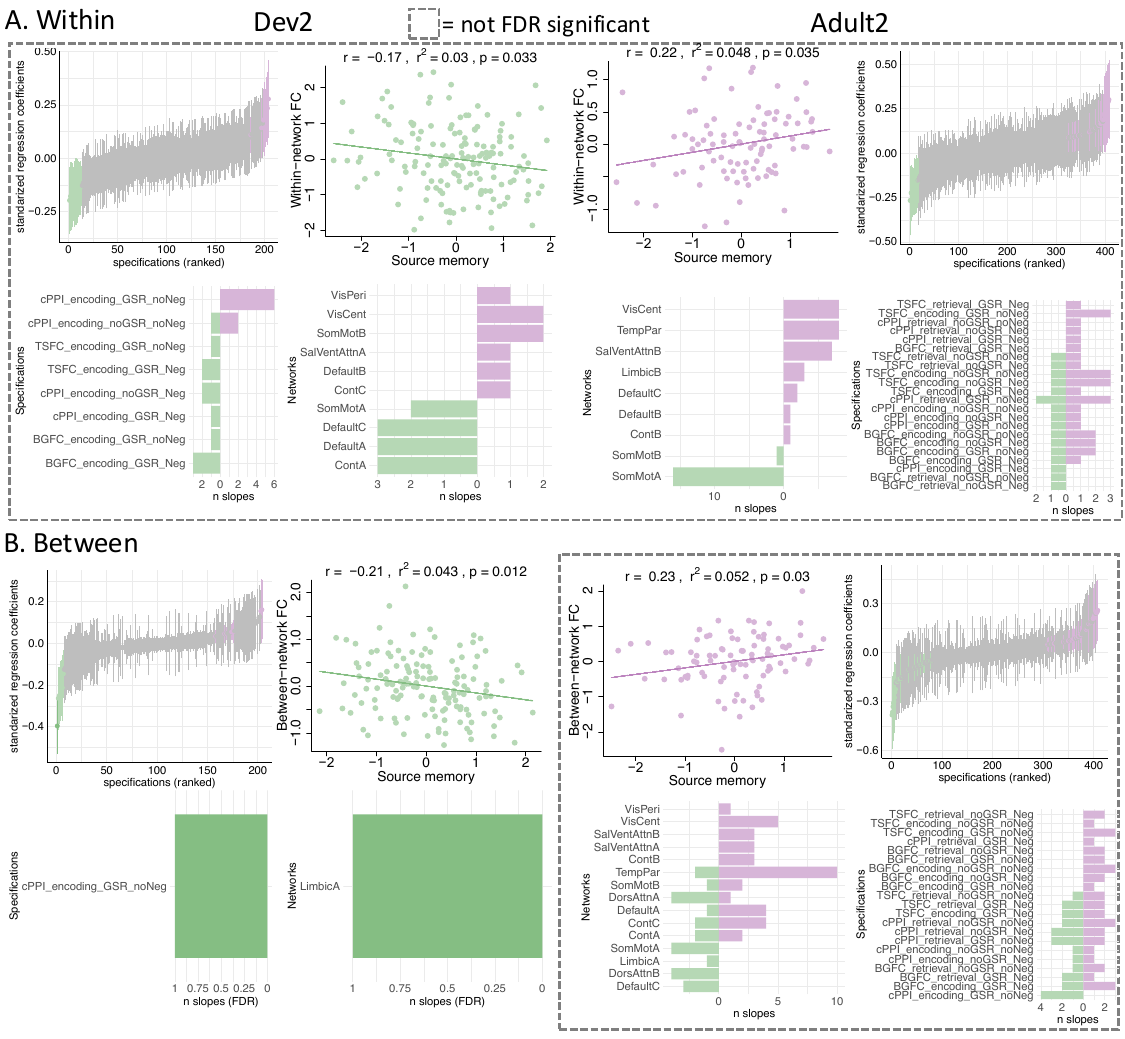

#### Figure 10. Task-general FC and memory performance – Dev2 & Adult2

Results shown for (**A**) within-network task-general FC (TGFC) for Dev1 and Adult1, (**B**) between-network TGFC for Dev2 and Adult2. Specification curves for associations with age (first and third row, first and fourth columns). Example scatter plots (first and third row, second and third columns) of median regression coefficient with uncorrected P values < 0.05 (FC and age residualized by model covariates for visualization). Involved analytic approaches (second and fourth row, first and fourth columns) and networks (second and fourth row, second and third columns). TSFC=Task-state FC. BGFC=Background FC.

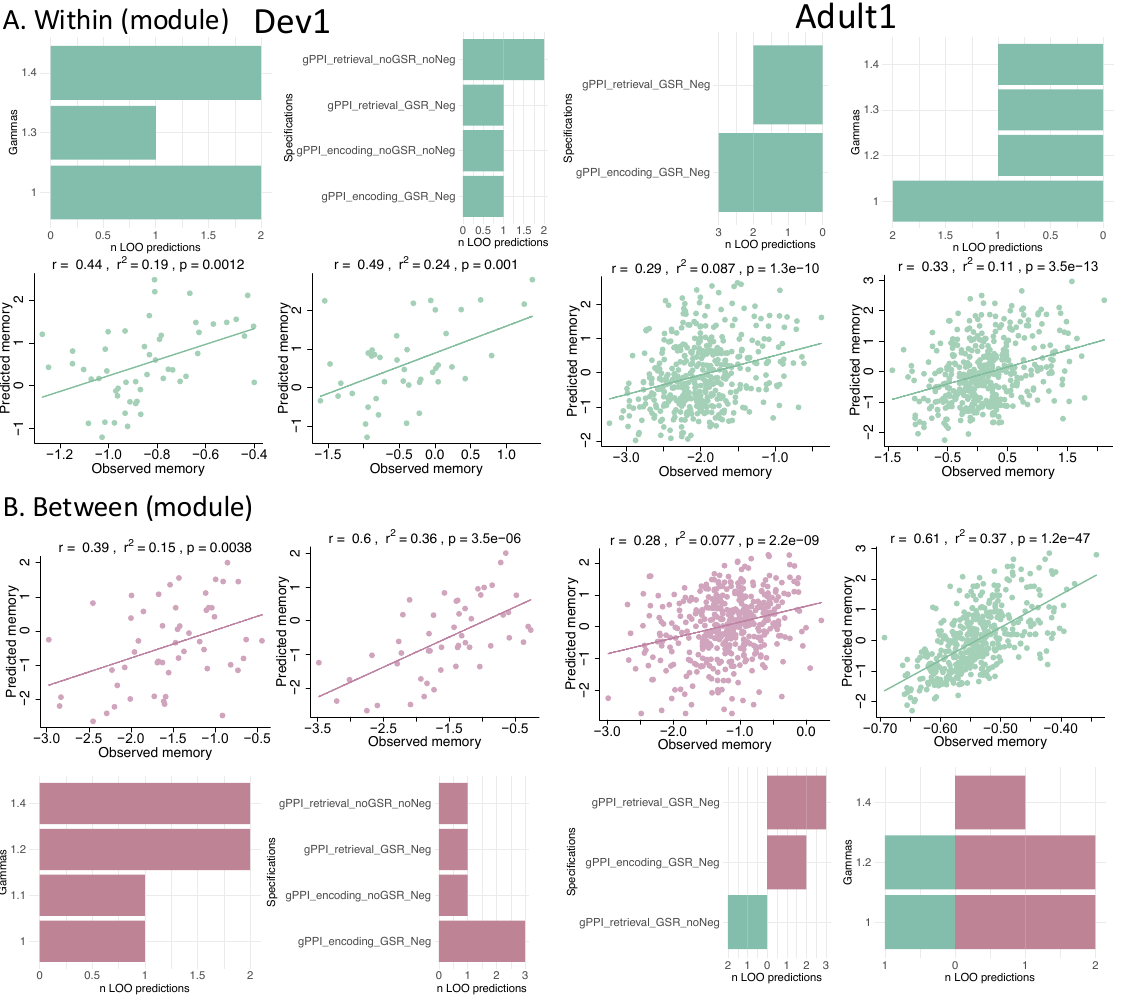

#### Figure 11. Memory-modulated FC and memory prediction Dev1 and Adult1

Within-network, memory-modulated FC in (**A)** Adult1 and **(B)** Dev1: significant networks (first column; P_perm_ < 0.05), corresponding specifications (second column), and, in the network showing the strongest associations, the scatter plot between observed memory performance and predicted memory performance based on FC.

Greenish: original FC-memory association negative, “redish: original FC-memory association positive.

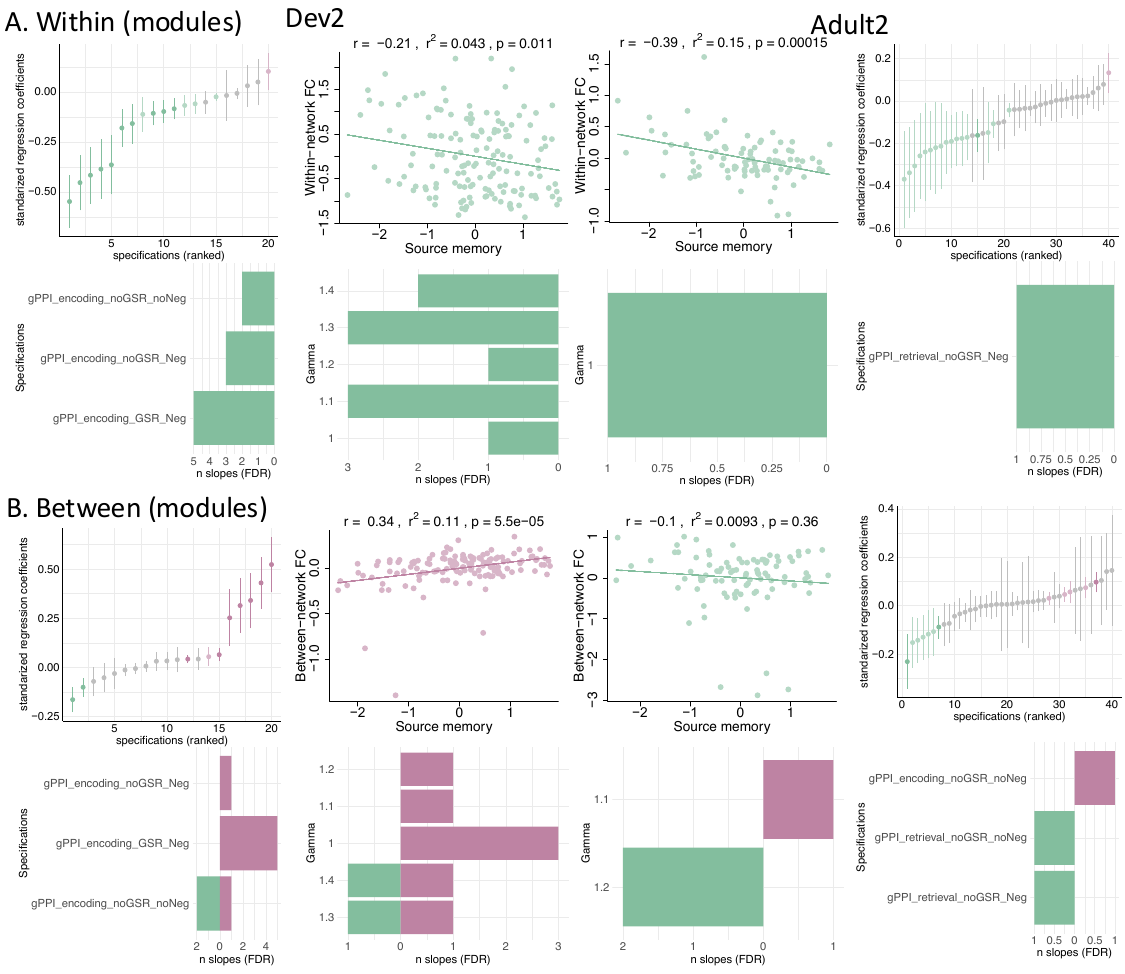

#### Figure 12. Memory-modulated FC in modularity-derived networks – Dev2 & Adult2

Results shown for (**A**) within-network memory-modulated FC (MMFC) for Dev1 and Adult1, (**B**) between-network MMFC for Dev2 and Adult2. Specification curves for associations with age (first and third row, first and fourth columns). Example scatter plots (first and third row, second and third columns) of median regression coefficient with uncorrected P values < 0.05 (FC and age residualized by model covariates for visualization). Involved analytic approaches (second and fourth row, first and fourth columns) and network resolution parameters γ (lower yields fewer networks, second and fourth row, second and third columns).

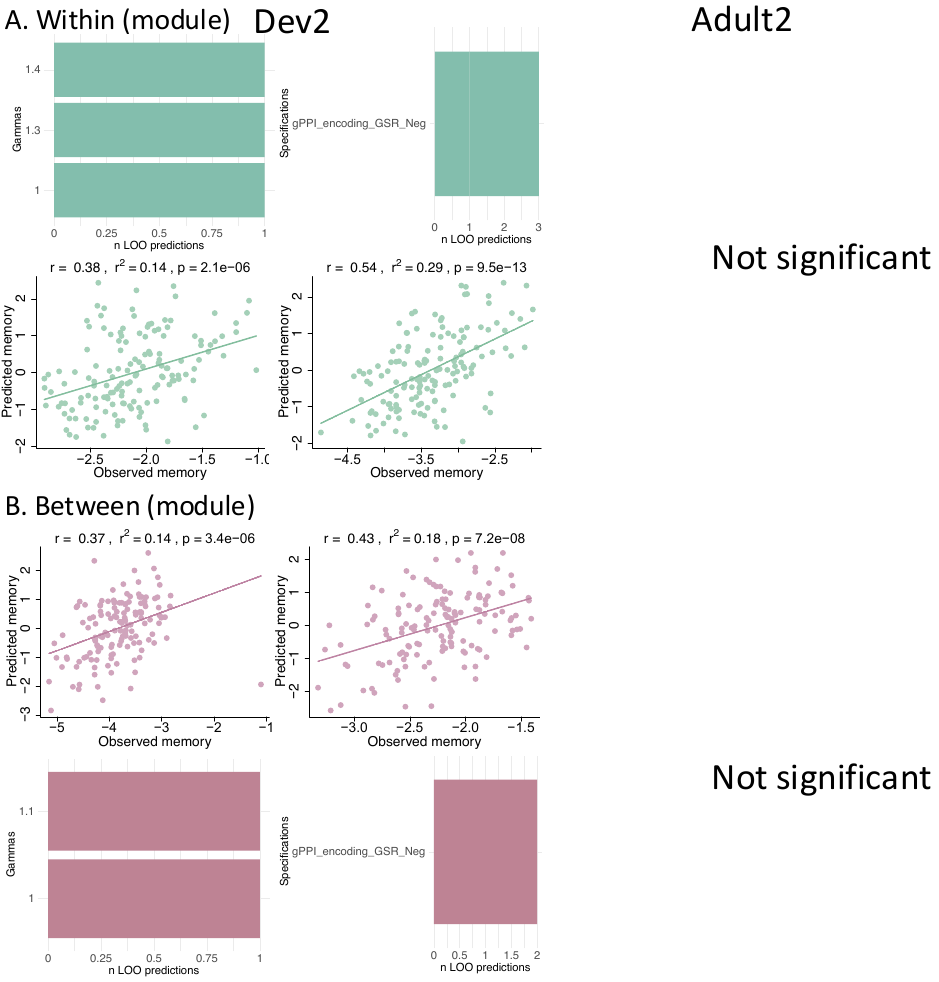

#### Figure 13. Memory-modulated FC and memory prediction Dev2 and Adult2

**A**. Within- and (**B)** between-network, task-general FC and memory predictions in Dev2 and Adult2: significant gamma resolution (first row, first column; P_perm_ < 0.05), corresponding specifications (first row, second column), and, in the network showing the weakest (first and third column) and strongest (second and fourth column) associations (second and fourth rows), the scatter plot between observed memory performance and predicted memory performance based on FC.
